# CRISPR targeting of MEIOTIC-TOPOISOMERASE VIB-dCas9 to a recombination hotspot is insufficient to increase crossover frequency in Arabidopsis

**DOI:** 10.1101/2021.02.01.429210

**Authors:** Nataliya E. Yelina, Sabrina Gonzalez-Jorge, Dominique Hirsz, Ziyi Yang, Ian R. Henderson

## Abstract

During meiosis, homologous chromosomes pair and recombine, which can result in reciprocal crossovers that increase genetic diversity. Crossovers are unevenly distributed along eukaryote chromosomes and show repression in heterochromatin and the centromeres. Within the chromosome arms crossovers are often concentrated in hotspots, which are typically in the kilobase range. The uneven distribution of crossovers along chromosomes, together with their low number per meiosis, creates a limitation during crop breeding, where recombination can be beneficial. Therefore, targeting crossovers to specific genome locations has the potential to accelerate crop improvement. In plants, meiotic crossovers are initiated by DNA double strand breaks (DSBs) that are catalysed by SPO11 complexes, which consist of two catalytic (SPO11-1 and SPO11-2) and two non-catalytic subunits (MTOPVIB). We used the model plant *Arabidopsis thaliana* to target a dCas9-MTOPVIB fusion protein to the *3a* crossover hotspot via CRISPR. We observed that this was insufficient to significantly change meiotic crossover frequency or pattern within *3a*. We discuss the implications of our findings for targeting meiotic recombination within plant genomes.

## Introduction

Meiosis is a specialized eukaryotic cell division where a single round of DNA replication and two rounds of chromosome segregation result in haploid gametes required for sexual reproduction (Mercier et al. 2015; Villeneuve and Hillers 2001). During prophase I of meiosis, homologous chromosomes undergo programmed recombination, which can result in reciprocal crossover (Mercier et al. 2015; Villeneuve and Hillers 2001). Crossovers contribute to genetic variation in progeny and result in new haplotypes, which can allow combination of useful traits in crop species (Taagen et al. 2020). However, recombination frequency and pattern can significantly limit breeding, as crossovers are relatively low per meiosis (typically 1-2 per chromosome) and show a highly uneven distribution (Taagen et al. 2020; Mercier et al. 2015). For example, crossovers in wheat, barley and maize occur predominantly in the sub-telomeric regions (Darrier et al. 2017; Rodgers-Melnick et al. 2015; Higgins et al. 2012; Mascher et al. 2017), which can cause linkage drag in low-recombination regions that are under selection. Therefore, technology to increase global crossover numbers, or induce recombination at loci of choice, have the potential to substantially accelerate crop breeding.

Crossovers are initiated by double-strand breaks (DSBs) catalysed by the conserved transesterase SPO11 (Keeney et al. 1997; Bergerat et al. 1997). SPO11 is a homolog of the archaeal topoisomerase VI catalytic A subunit that acts with non-catalytic B subunits in A_2_B_2_ heterodimers (Robert et al. 2016; Bouuaert and Keeney 2016). In Arabidopsis, two non-redundant homologs of the topoisomerase VI A subunit, SPO11-1 and SPO11-2, are required to generate meiotic DSBs (Stacey et al. 2006; Grelon et al. 2001; Hartung et al. 2007). The meiotic topoisomerase VIB-like subunits, MTOPVIB, interact with both SPO11-1 and SPO11-2 to catalyse meiotic DSBs in Arabidopsis and rice (Bouuaert and Keeney 2016; Vrielynck et al. 2016; Fu et al. 2016). During catalysis, SPO11 becomes covalently bound to DNA and is then removed bound to a short oligonucleotide, via endonuclease activities (Neale et al. 2005; Choi et al. 2018). The resulting DSB 5’-end is then digested by exonucleases to produce 3’ overhanging single-strand DNA (ssDNA) at each end of the DSB (Hunter 2015). Meiotic ssDNA associates with the recombinases RAD51 and DMC1 to promote ssDNA strand invasion of a homologous chromosome or a sister chromatid (Hunter 2015). Invasion of homologous DNA generates a displacement loop (D-loop), which allows extension of the 3’ ssDNA via DNA synthesis using the homologous DNA sequence as a template (Hunter 2015).

Following interhomolog or intersister strand invasion, alternative DNA repair pathways are followed during meiosis (Hunter 2015). First, the D-loop may be disassociated from the invaded template and returned to the parental chromosomes, where it is repaired as a non-crossover (Hunter 2015). If DNA synthesis occurred over a polymorphic site following inter-homolog strand invasion this may result in a gene conversion (Hunter 2015). In plants, non-crossover repair is promoted via the activity of several non-redundant proteins that include the FANCM, RECQ4A and RECQ4B helicases, FIGL1 and FLIP1 (Crismani et al. 2012; Girard et al. 2015; Fernandes et al. 2018; Séguéla-Arnaud et al. 2015). Alternatively, capture of the second resected 3’ end, followed by DNA synthesis, can form a double Holliday junction joint molecule (dHJ-JM) (Hunter 2015). The Class I pathway acts to stabilize dHJs and promotes their resolution as a crossover (Börner et al. 2004; Chelysheva et al. 2012, 2007; Higgins et al. 2008, 2004; Macaisne et al. 2011, 2008; Wijeratne et al. 2006; Manhart and Alani 2016; Jackson et al. 2006). In Arabidopsis, an estimated ~150-250 DSBs mature into ~10 crossovers per meiosis, with the remaining DSBs repaired as non-crossovers (Rowan et al. 2019; Ferdous et al. 2012; Wijnker et al. 2013). This indicates that the anti-crossover pathways mediate repair of the majority of meiotic DSBs as non-crossovers.

Chromosome structure, chromatin and epigenetic information also exert a significant influence on meiotic recombination. At the fine-scale, meiotic DSBs and crossovers tend to cluster in narrow (kilobase) regions called hotspots (Choi and Henderson 2015). In plants and budding yeast, meiotic DSB hotspots frequently occur in nucleosome-depleted regions associated with gene control regions (He et al. 2017; Pan et al. 2011; Choi et al. 2018). Furthermore, RNA-directed DNA methylation and elevated nucleosome occupancy are sufficient to suppress crossovers within an Arabidopsis recombination hotspot (Yelina et al. 2015). Meiotic DSB formation and repair occur in the context of proteinaceous chromosome axis, which underpins meiotic chromosome architecture (Zickler and Kleckner 1999). Sister chromatids are organized as linear arrays of chromatin loops connected to the axis (Zickler and Kleckner 1999). In plants, the chromosome axis includes the HORMA domain protein ASY1 (a homolog of yeast Hop1) and its interacting partners ASY3 and ASY4, which promote DMC1-mediated interhomolog synapsis and recombination (Armstrong et al. 2002; Sanchez-Moran et al. 2007; Chambon et al. 2018; Ferdous et al. 2012). The axis also includes cohesin complexes containing the meiosis-specific REC8 α-kleisin subunit, which coheres sister chromatids and anchors the chromatin loops to the axis (Cai et al. 2003; Chelysheva et al. 2005). As prophase I progresses, the chromosomes synapse and the synaptonemal complex is installed between them, coincident with crossover maturation (Zickler and Kleckner 1999; Hunter 2015).

Work in budding yeast has shown that tethering SPO11, or its interacting partners, using DNA binding domains is sufficient to create recombination hotspots *de novo* (Acquaviva et al. 2013; Pecina et al. 2002). In recent years, several technologies have emerged with the potential to tether factors of interest to specific loci. For example, translational fusions of SPO11 with zinc finger domains, TAL repeats and dCas9 have been used to target meiotic DSBs to loci of choice in budding yeast (Sarno et al. 2017). In this study, we targeted an MTOPVIB-dCas9 fusion protein to the previously characterized *3a* crossover hotspot in *Arabidopsis thaliana*. The catalytically <u>d</u>ead *Streptococcus pyogenes* Cas9 (<u>d</u>Cas9) carries two amino acid substitutions (D10A and H841A) that abolish its endonuclease activity, but do not impair its ability to bind target DNA via guide RNAs (Qi et al. 2013). We used high-resolution crossover mapping to determine *3a* recombination frequency and distribution with and without MTOPVIB-dCas9 targeting. We did not observe significant changes to crossover frequency or pattern with the *3a* hotspot compared to wild type. This indicates that recruitment of the DSB machinery to Arabidopsis hotspots is insufficient to change crossover recombination.

## Results

### *MTOPVIB-dCas9* functionally complements *mtopvib*

Meiotic DSBs catalysed by Arabidopsis SPO11-1, SPO11-2 and MTOPVIB are essential to initiate crossover formation (Fig. 1A) (Grelon et al. 2001; Vrielynck et al. 2016; Stacey et al. 2006; Hartung et al. 2007). We translationally fused *Streptococcus pyogenes* dCas9 to the C-terminus of Arabidopsis MTOPVIB and asked whether the fusion protein complements the function of wild type *MTOPVIB* (Fig. 1B-F). We expressed an *MTOPVIB-dCas9* translational fusion gene under the control of the endogenous *MTOPVIB* promoter and terminator, in an *mtopvib-2* (hereafter, *mtopvib*) null mutant background (Vrielynck et al. 2016). Crossovers physically link homologous chromosomes during prophase I of meiosis ensuring balanced chromosome segregation. Therefore, an absence of meiotic DSBs and crossovers in *mtopvib* (or *spo11-1* and *spo11-2*) mutants leads to unbalanced, aneuploid gametes and almost complete sterility (Fig. 1B-1C) (Grelon et al. 2001; Vrielynck et al. 2016; Stacey et al. 2006; Hartung et al. 2007). For example, we observed an average of ~1.9 +/− 1.7 seeds per fruit (silique) in *mtopvib*, compared to ~64.5 +/− 8.2 in the wild type (Two-tailed t-test, *P*<0.00001) (Fig. 1B-1C and Supplemental Table 1). In contrast, *MTOPVIB-dCas9 mtopvib* shows an average seed set of 66.1 +/−3.3 seeds per silique that was not significantly different from wild type (Two-tailed t-test, *P*=0.63) (Fig. 1B-1C, Supplemental Table 1), indicating that the MTOPVIB-dCas9 fusion protein functionally complements *mtopvib*. To further confirm this, we used fluorescent crossover reporters to measure genetic distances (crossover frequency) in two intervals; *CTL2.10*, an interstitial region on chromosome 2, and *CTL5.1*, a sub-telomeric region on chromosome 5 (Wu et al. 2015), in *MTOPVIB-dCas9 mtopvib* and wild type (Figure 1D-1F and Supplemental Tables 2 and 3). We found that mean genetic distances in these intervals were not significantly different between wild type and *MTOPVIB-dCas9 mtopvib* (Whitney-Mann tests, *P*=0.54 and 0.68, respectively). This further demonstrates that the MTOPVIB-dCas9 fusion protein is functional and supports a normal level of crossover.

**Figure 1.**
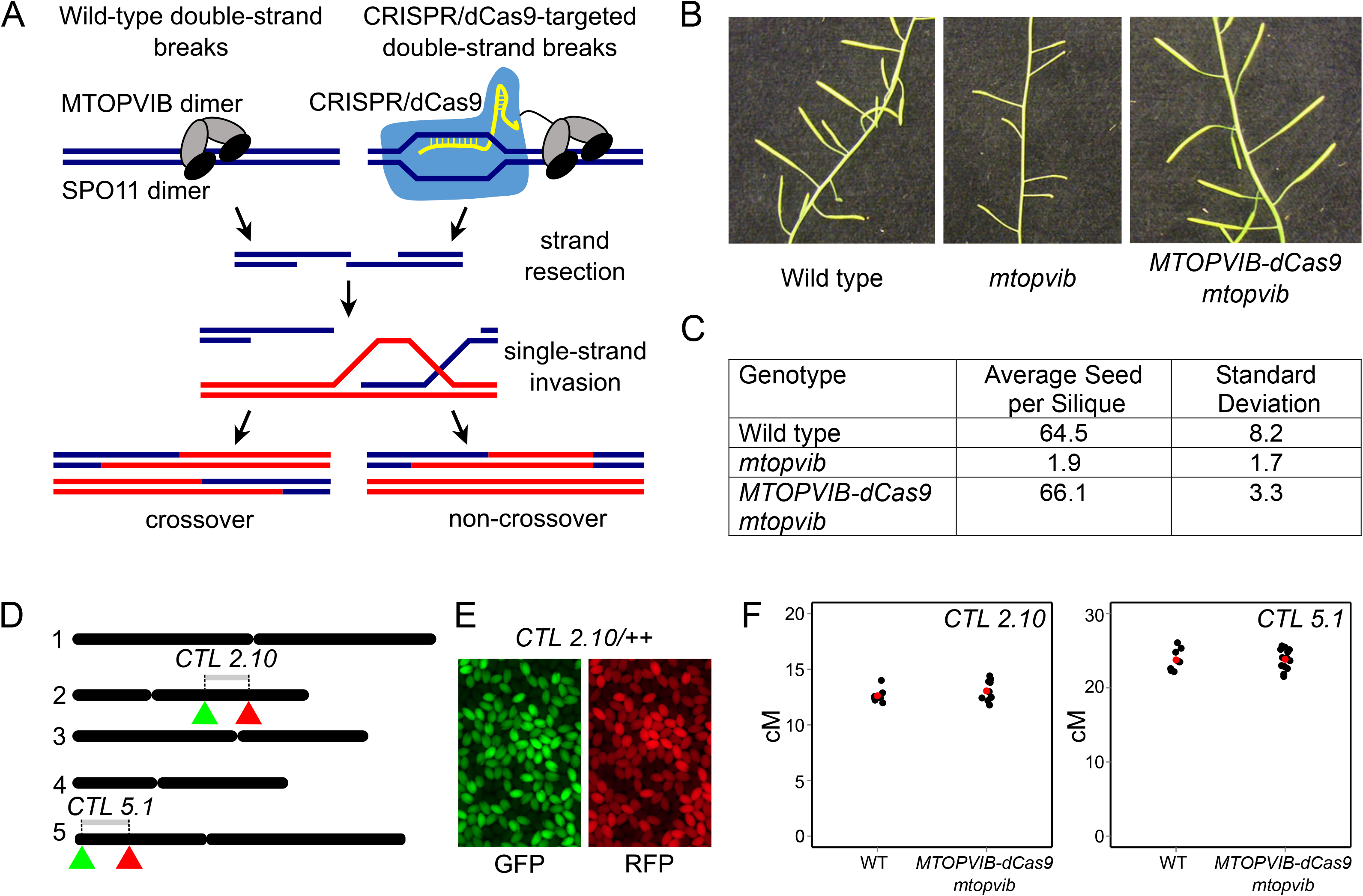
Complementation of Arabidopsis *mtopvib* with MTOPVIB fused to catalytically inactive Cas9 (MTOPVIB-dCas9). **(A)** Wild type and synthetic pathways to generate meiotic double-strand breaks. Homologous chromosomes shown as red and blue lines, MTOPVIB as grey ovals, SPO11 homologues as black ovals, CRISPR/dCas9 shown in blue, guide RNA paired to a genomic locus in yellow. **(B)** Arabidopsis inflorescences showing long fruit (siliques) in wild type and complementing lines (MTOPVIB-dCas9 in *mtopvib* background) and short fruit (siliques) in *mtopvib*. **(C)** Average seed count per silique and standard deviation for each genotype. **(D)** Seed-based reporter systems to measure crossovers in two tester intervals, interstitial *CTL 2.10* on chromosome 2 and sub-telomeric *CTL 5.1* on chromosome 5. Five Arabidopsis chromosomes are shown as black lines, reporter transgenes, *eGFP* and *dsRED*, represented by green and red triangles, respectively. (**E)** Fluorescent micrographs showing *CTL 2.10* (*GFP RFP/++)* seed using green or red fluorescent filters. **(F)** Genetic distances of *CTL 2.10* and *CTL 5.1* in wild type and *MTOPVIB-dCas9 mtopvib*. Each black dot represents crossover frequency in an individual plant, red dots denote mean crossover frequencies. Whitney-Mann test showed that mean crossover frequencies in *CTL 2.10* and *CTL 5.1* were not significantly different between wild type and complementing lines (*P* values of 0.54 and 0.68, respectively).

### Targeting MTOPVIB-dCas9 to the *3a* meiotic crossover hotspot

We chose to target MTOPVIB-dCas9 to the *3a* crossover hotspot (Yelina et al. 2012, 2015; Choi et al. 2013), which is located in a sub-telomeric region of chromosome 3 (Fig. 2A, 2B and Supplemental Table 4). *3a* is a 5.8 kb region with a genetic distance of ~0.2 cM (33.3 cM/Mb) in F_1_ hybrids between Col-0 (hereafter, Col) and Ler-0 (hereafter, Ler) *Arabidopsis thaliana* accessions (Yelina et al. 2012, 2015; Choi et al. 2013). Crossover rates within *3a* are up to ~17-fold higher than the chromosome 3 average of 4.77 cM/Mb in male meiosis (Giraut et al. 2011). We chose to target the *3a* hotspot first because data from budding yeast showed that tethering SPO11 to recombination hotspots leads to additional DSB formation (Sarno et al. 2017), whereas tethering to recombination ‘cold’ regions exhibited variable and less predictable stimulations (Robine et al. 2007; Ito et al. 2014; Panizza et al. 2011; Sarno et al. 2017; Pan et al. 2011). Second, *3a* crossover levels are below their potential maximum in wild type, as we have previously shown a ~40% increase in *3a* crossover frequency in *met1* mutants (Yelina et al. 2012). Third, we have an established ‘pollen typing’ assay that allows us to measure *3a* crossover rates and fine-map crossover positions in this region (Yelina et al. 2012, 2015; Choi et al. 2013). We designed guide RNAs (gRNAs) to target three regions within *3a*: (i) the At3g02880 promoter and 5’ end (hereafter, *3a-P*), (ii) the At3g02880 gene body (hereafter, *3a-B*), and (iii) the intergenic region between At3g02880 and At3g02885 (hereafter, *3a-I*) (Fig. 2A, 2B and Supplemental Tables 4–5). Notably these regions vary in nucleosome occupancy, which is a major determinant of meiotic DSB levels in Arabidopsis (Fig. 2A) (Choi et al. 2018).

**Figure 2.**
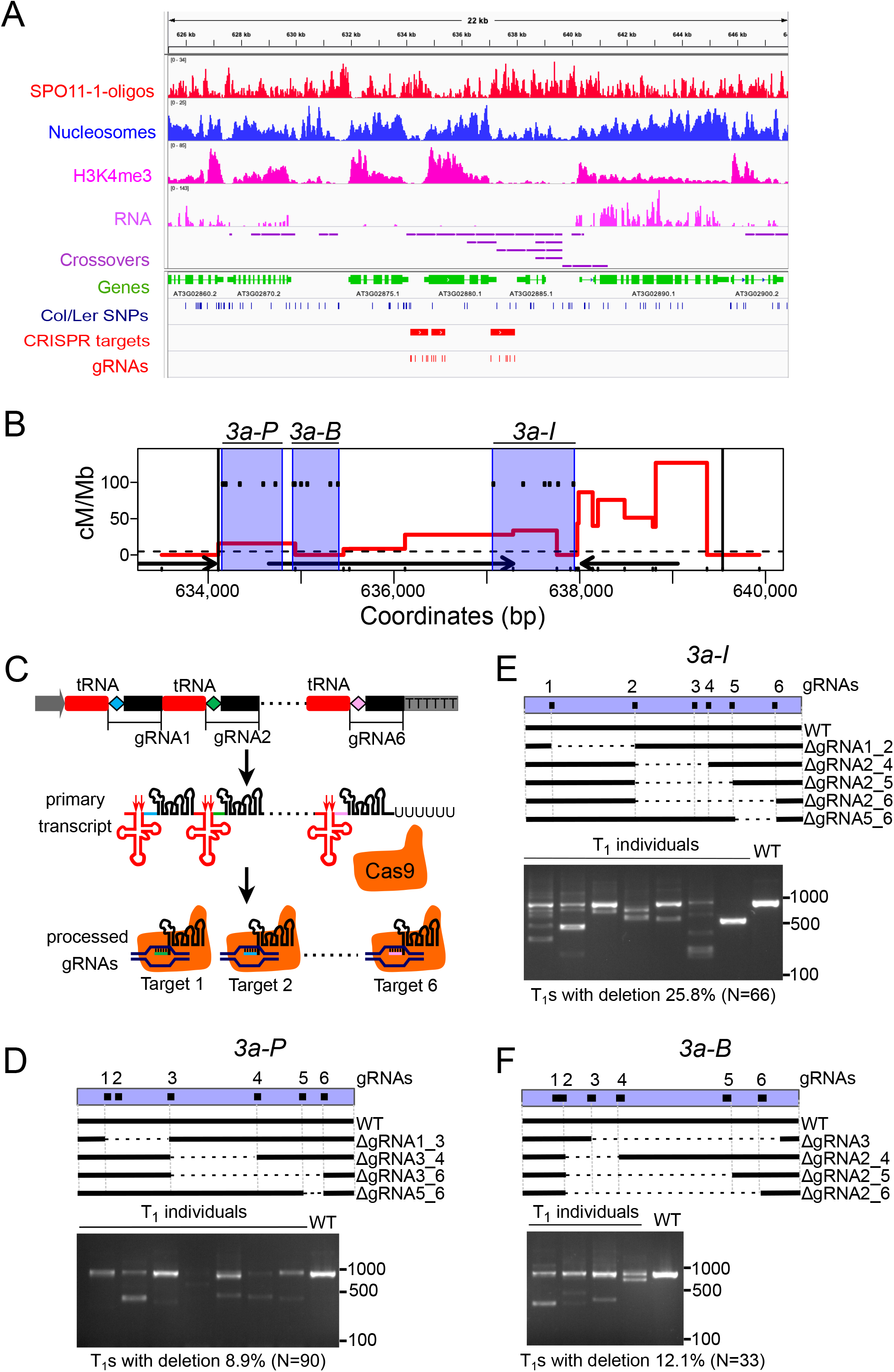
Testing gRNAs targeting *3a* meiotic recombination hotspot via a catalytically active Cas9. **(A)** Histograms for the chromosome 3 sub-telomeric region showing library size normalized coverage values for SPO11-1-oligonucleotides (red), nucleosome occupancy (blue, MNase-seq), H3K4me3 (pink, ChIP-seq), RNA-seq (lilac) and crossovers (purple). Positions of CRISPR target regions are shown as red rectangles and individual gRNA target loci as red ticks. TAIR 10 gene annotations are shown in green and single nucleotide polymorphisms between Col and Ler as blue ticks. **(B)** *3a* crossover profile, red line (centimorgans per megabase, cM/Mb), in Col/Ws *MTOPVIB-dCas9 mtopvib* F_1_s. Black vertical lines delineate borders of the *3a* hotspot, ticks on the *x*-axis represent polymorphisms between Col and Ws. Black arrows represent genes, dashed horizontal line - male chromosome 3 average crossover frequency. Six gRNAs were designed to target each of the three regions within *3a*, *3a-P*, *3a-B* and *3a-I*, shaded in blue. gRNA target sites are shown as black ticks within the blue shaded areas. **(C)** Multiplexing six gRNAs via endogenous tRNA-processing system. Schematic representation of a gRNA-tRNA transgene containing tandemly arranged tRNAs and gRNAs. Pol III promoter - grey arrow, terminator - grey rectangle, guide RNA-specific spacers are shown as diamonds of different colour (blue, green or pink), conserved gRNA scaffold shown as black rectangles, tRNA as red rectangles. The primary transcript is cleaved by endogenous RNase P and RNase Z (red arrows) to release mature tRNA (red cloverleaf structure). Processed mature gRNAs guide catalytically active Cas9 (orange) to specific targets. gRNAs 3-5 and their targets are not shown. **(D)** CRISPR/Cas9-induced deletions in *3a-P*. *3a-P* is shown as blue rectangle, six gRNAs as black squares. Wild-type and deleted regions within *3a-P* are shown by black and dashed lines, respectively. Midori-green-stained agarose gel image shows PCR-amplified *3a-P* in wild type (WT) and representative individual T_1_s. Lower than wild type molecular weight products result from CRISRP/Cas9-mediated deletions in *3a-P*. Percentage of T_1_s with CRISRP/Cas9 induced deletions and the total number of T_1_s analysed are indicated under the agarose gel image. **(E)** As in **(D)** but for *3a-I* region. **(F)** As in **(D)** but for *3a-B* region.

### Testing gRNA gene editing efficiency using catalytically active Cas9

We designed a total of 18 gRNAs within *3a*, six targeting each of the three regions within *3a* (*3a-P*, *3a-B* and *3a-I*), with the rationale that multiple gRNAs may increase the efficiency of targeting, compared to a single gRNA (Fig. 2A-2F and Supplemental Table 5) (Chavez et al. 2016; Sarno et al. 2017). To simultaneously express six gRNAs using one T-DNA construct, we used an approach successfully employed in Arabidopsis, rice and wheat, where multiple gRNAs are expressed as part of a tRNA-gRNA synthetic transcript (Xie et al. 2015; Hui et al. 2019; Wang et al. 2018). We designed and assembled six tandemly arranged pre-tRNA-gRNA modules differing only in the sequences of gRNA spacers (Fig. 2C). pre-tRNA-gRNA synthetic transcripts mimic native tRNA-snoRNA43 transcripts in plants, allowing RNase P and Z to cleave the tRNA structure and release mature gRNAs (Fig. 2C) (Phizicky and Hopper 2010; Xie et al. 2015). We tested the efficiencies of *in silico* designed gRNAs by co-expressing 6ˣ(pre-tRNA-gRNA) cassettes targeting *3a-P*, *3a-B* or *3a-I* with catalytically active *S. pyogenes* Cas9 in wild type Col (Fig. 2D-2F) (Schiml et al. 2016). We transformed *Cas9-gRNA-P*, *Cas9-gRNA-B* and *Cas9-gRNA-I* constructs into Arabidopsis and analysed gene editing events within *3a* in T_1_ progeny. T_1_ individuals are usually chimeric due to somatic gene editing events (Hui et al. 2019). Using PCR amplification across the gRNA target sites, we observed deletions in the respective target regions in 8.9, 12.1 and 25.8% of T_1_ progeny of *Cas9-gRNA-P*, *Cas9-gRNA-B* and *Cas9-gRNA-I*-transformed plants (Fig. 2D-2F and Supplemental Table 6). Sanger sequencing of these PCR products confirmed deletions associated with 17 of the 18 tested gRNAs (Fig. 2D-2F and Supplemental Figures S1, S2, and Supplemental Table 6).

In addition, we generated a synthetic 6ˣ(pre-tRNA-gRNA) construct to express previously reported gRNAs targeting 6 Arabidopsis genes (At1g69320, At1g26600, At2g27250, At3g27920, At4g25530 and At5g35620) outside *3a* to use as a negative control (Yamaguchi et al. 2017; Hahn et al. 2017; Pyott et al. 2016; Gallego-Bartolomé et al. 2018). We refer to this construct as *Cas9*-*non-3a-gRNA*. We transformed *Cas9*-*non-3a-gRNA* into wild type Col and observed gene editing events in the target genes in ~4-50% of the T_1_ progeny (Supplemental Figure S3 and Supplemental Table 7). In summary, we obtained a set of gRNAs robustly targeting the Arabidopsis genome within and outside the *3a* crossover hotspot.

### Analysis of *3a* crossovers in the presence of MTOPVIB-dCas9 and gRNAs

We next asked whether combining *MTOPVIB-dCas9* and gRNAs that target *3a-P*, *3a-B* or *3a-I* would affect 3a crossover rates or distribution. Crossover detection at *3a* hotspot relies on the segregation of DNA sequence polymorphisms through meiosis (Yelina et al. 2012, 2015; Choi et al. 2013). As *MTOPVIB-dCas9 mtopvib* lines were in the Col background, we generated transgenic lines expressing gRNAs in a different Arabidopsis accession, Ws-4 (hereafter, Ws), that was also heterozygous for a *mtopvib* mutation (*mtopvib-1*) (Vrielynck et al. 2016). The resulting lines, each of which carried a 6ˣ(pre-tRNA-gRNA) transgene targeting *3a-P*, *3a-B* or *3a-I*, or six Arabidopsis genes outside *3a*, were called *gRNA-P*, *gRNA-B*, *gRNA-I* or *non-3a gRNA*. Ws had a single nucleotide polymorphism (SNP) in position −3 relative to PAM in a target site of one of the *gRNA-B*-specific gRNAs, the remaining 17 *3a*-specific gRNAs we used targeted regions without any polymorphisms between Col and Ws. We crossed *MTOPVIB-dCas9 mtopvib* in the Col background to *gRNA-P*, *gRNA-B* and *gRNA-I* lines in the Ws background. We then identified F_1_ progeny that were *mtopvib* null mutants and that expressed both the *MTOPVIB-dCas9* and gRNA transgenes (Fig. 3A, Supplemental Figure S4). We also crossed Col *MTOPVIB-dCas9 mtopvib* to Ws *MTOPVIB/mtopvib* to generate a ‘no gRNA’ F_1_ population as a negative control.

**Figure 3.**
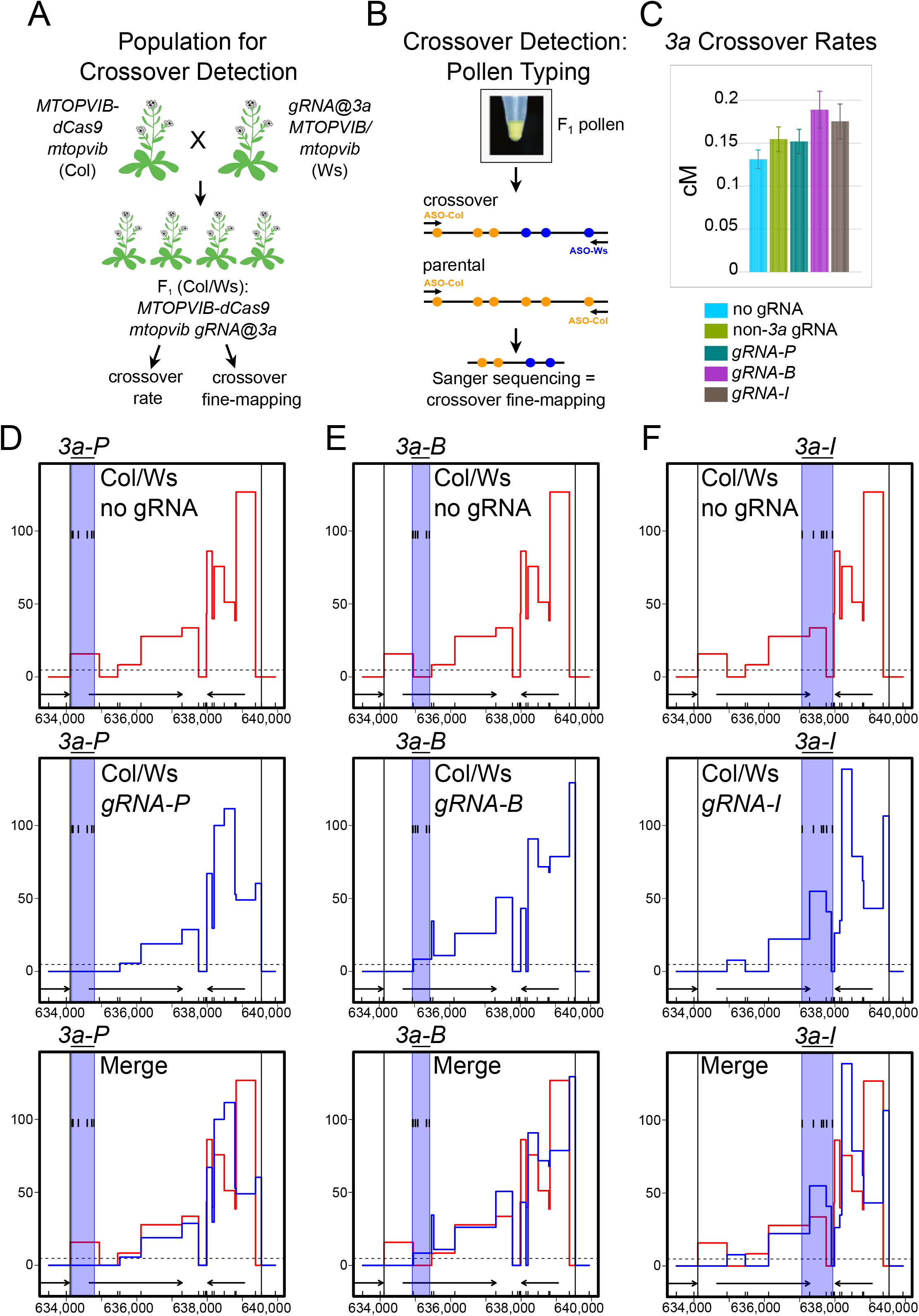
*3a* crossover rates in targeted and wild type F_1_ hybrids. **(A)** Generation of F_1_ populations for fine-scale crossover analysis via ‘pollen typing’. Col *mtopvib* lines complemented with *MTOPVIB-dCas9* were crossed to Ws *MTOPVIB/mtopvib* carrying a guide RNA transgene targeting *3a* crossover hotspot (gRNA@3a). The resulting Col/Ws F_1_ populations were selected for the presence of *MTOPVIB-dCas9* transgene, absence of wild-type *MTOPVIB* (*mtopvib*) and presence of gRNA transgene. A similar crossing scheme was performed for negative controls, ‘no gRNA’ and ‘non-3a gRNA’, not shown. **(B)** Schematic representation of ‘pollen typing’. Genomic DNA extracted from F_1_ pollen is subject to PCR amplification with allele specific oligonucleotides (ASO) to determine the concentration of recombinant crossover molecules relative to parentals. Recombinant molecules are then subject to Sanger sequencing to determine crossover distribution within the hotspot. DNA molecules shown as black lines. Yellow and blue circles represent Col- and Ws-specific polymorphisms, respectively. **(C)** *3a* crossover frequencies in centimorgans (cM) measured by pollen typing in five different F_1_ populations. Error bars represent standard deviation. **(D)** Fine-scale *3a* crossover profiles in the presence or absence of *gRNA-P* gRNAs targeting *3a-P*. *3a* recombination rates in centimorgans per megabase (cM/Mb) were analysed by pollen typing. Black vertical lines delineate borders of *3a* hotspot, ticks on the *x*-axis represent polymorphisms between Col and Ws. Black arrows represent genes, dashed horizontal line - male chromosome 3 average crossover frequency. Blue shaded area (*3a-P*) marks guide RNA target region with black ticks representing individual guide RNA target sites. Recombination rates in Col/Ws *MTOPVIB-dCas9 mtopvib* F_1_s in the absence of guide RNAs are shown in red and in the presence of *gRNA-P* gRNAs - in blue. **(E)** As in **(D)**, but for *3a-B*. **(F)** As in **(D)** but for *3a-I*.

Given the *3a* crossover rate of ~0.2 cM, to characterise ~100 crossover events, it is necessary to assay ~50,000 meioses. To achieve this we employed ‘pollen typing’, which is a PCR-based assay used to amplify and quantify crossover and parental molecules from pollen DNA (Fig. 3B) (Drouaud and Mézard 2011; Choi et al. 2017). To perform pollen typing, we first extract genomic DNA from F_1_ pollen. The pollen DNA contains *3a* parental and crossover molecules distinguishable by DNA sequence polymorphisms between the accessions (Col and Ws) (Fig. 3B). We perform two rounds of allele-specific PCR, using primers that anneal to polymorphic sites, to specifically amplify crossover or parental molecules (Fig. 3B). For quantification, we use titration where pollen template DNA is diluted until approximately half of PCR amplification reactions are negative (Drouaud and Mézard 2011; Choi et al. 2017). We also Sanger sequenced the amplified crossover molecules to map internal crossover locations within the *3a* hotspot (Drouaud and Mézard 2011; Choi et al. 2017).

We employed pollen typing to measure *3a* crossover frequency (genetic distance) and observed ~0.13-0.15 cM in Col/Ws F_1_s in the absence of gRNAs (Fig. 3C and Supplemental Table 8). We observed no significant crossover rate changes in F_1_ populations expressing *gRNA-B*, *gRNA-I* or *gRNA-P* (0.189 cM, chi-square test, *P*=0.44, 0.175 cM, *P*=0.64 and 0.152 cM, *P*=0.96, respectively), compared to negative controls (0.155 cM and 0.131 cM) (Fig. 3C and Supplemental Table 8). We Sanger sequenced between 77 and 90 crossover molecules for each F_1_ population and found that crossover profiles were very similar in the presence or absence of gRNAs targeting *3a* (Fig. 3D-3F and Supplemental Table 9). In all cases we observed lower crossover frequencies at the telomere-proximal end and higher crossover frequencies towards the centromere-proximal end of *3a* (Fig. 3D-3F and Supplemental Table 9). These data indicate that targeting MTOPVIB-dCas9 to *3a* does not have a strong effect on crossover rate or distribution.

In Arabidopsis, a minority of meiotic DSBs (~5-10%) are repaired as crossovers (Choi et al. 2018; Serrentino and Borde 2012; Copenhaver et al. 1998; Chelysheva et al. 2012, 2007; Giraut et al. 2011; Salomé et al. 2012). Non-crossovers are an alternative outcome of meiotic DSB repair and, therefore, we asked whether targeting MTOPVIB-dCas9 to *3a* could result in increased non-crossovers, measured via gene conversion. To detect gene conversion we used four F_2_ populations, which were the progeny of Col/Ws F_1_ expressing either *gRNA-P*, *gRNA-B* or *gRNA-I*, or ‘no gRNA’ as a negative control (Figure 4A). We employed a Kompetitive Allele-Specific PCR (KASP) assay to distinguish between SNP alleles (Figure 4B). We designed 12 KASP assays to distinguish between Col and Ws alleles within, as well as up to 5.2 kb up- and 4.5 kb downstream of *3a*. Physical distances between the markers used for KASP assays ranged from 0.7 to 2.0 kb, with an average of 1.3 kb (Supplemental Table 10). Initially, we used ~83-96 F_2_ individuals for each of the four F_2_ populations and detected 2 gene conversion events in the F_2_ population expressing *gRNA-P* and none in the other three populations, including the ‘no gRNA’ negative control. Next we increased *gRNA-P* and ‘no gRNA’ F_2_ population sizes to the total of ~470 individuals each but did not detect any additional gene conversion events (Fig. 4C-4E and Supplemental Table 11). Therefore, we did not observe any gene conversion events in *gRNA-B*, *gRNA-I* or the negative control, but observed two gene conversions out of 469 F_2_ individuals in *gRNA-P*. Next we performed a combination of Sanger sequencing and KASP assays at additional SNPs to confirm our initial results and to determine gene conversion tract lengths. We found that one of the gene conversion events, which had a Ws to Col to Ws genotype, occurred in a low polymorphism region and its tract length could vary from a minimum of 1 to a maximum of 1,763 bp. The other gene conversion event, which had a Col to Ws to Col genotype, occurred in a region more densely covered with polymorphisms. Its conversion tract could vary from a minimum of 729 to a maximum of 1,503 bp (Fig. 4E, Supplemental Table 12). Interestingly, both gene conversion events occurred 1.3-3 kb downstream of *gRNA-B* at adjacent, non-overlapping polymorphisms (Fig. 4E and Supplemental Tables 11-12).

**Figure 4.**
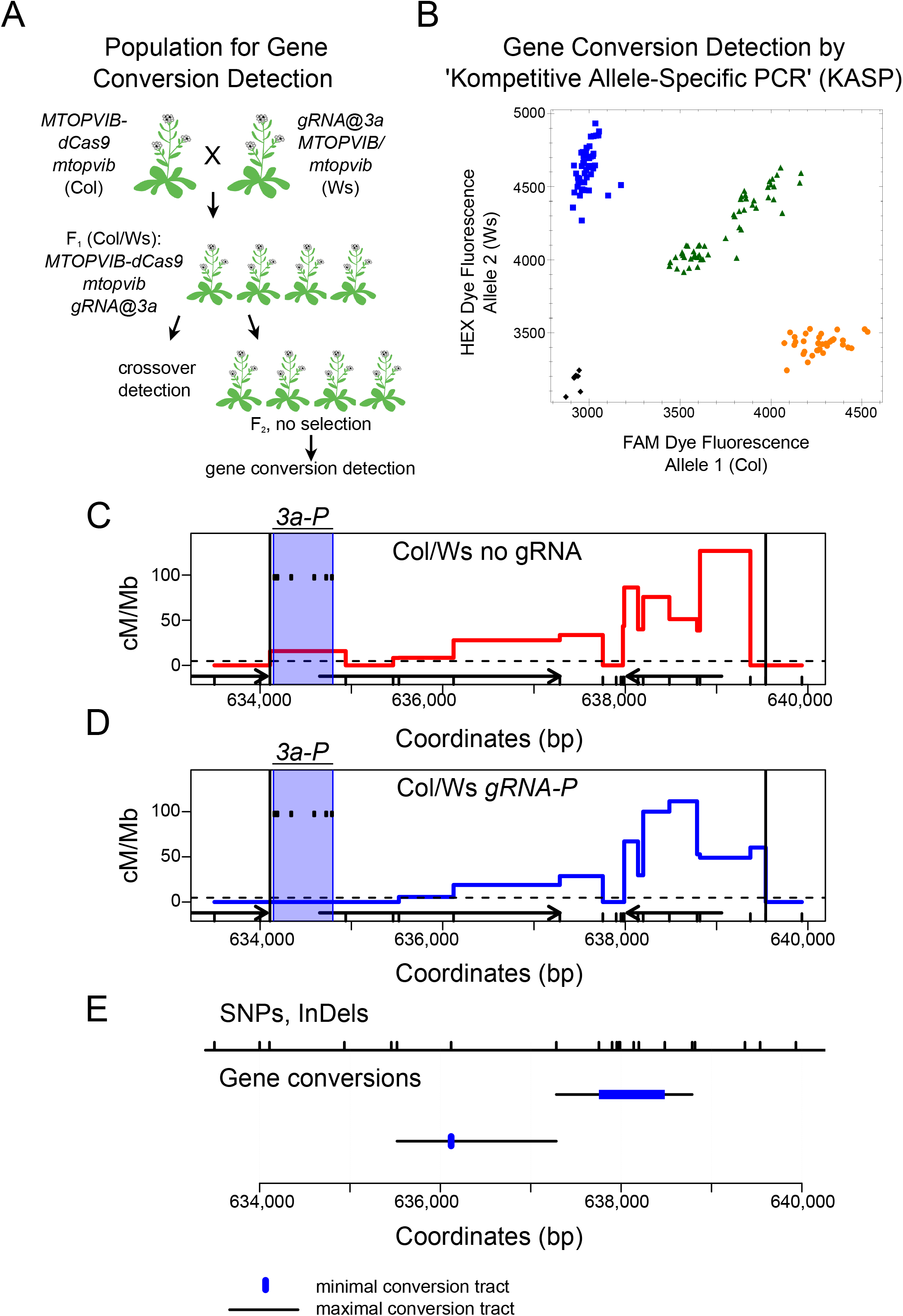
Gene conversions in *3a*. **(A)** Generation of F_2_ populations for gene conversion detection via Kompetitive Allele-Specific PCR (KASP). **(B)** An example plot showing allele discrimination via KASP assay for one single nucleotide polymorphism (SNP) between Col and Ws. Each dot represents an F_2_ individual. Different colours - yellow, blue, green and black - represent Col, Ws, heterozygous and a ‘no DNA’ control, respectively. **(C)** *3a* fine-scale crossover profile, red line in centimorgan per megabase (cM/Mb) in Col/Ws *MTOPVIB-dCas9 mtopvib* F_1_ population. Black vertical lines delineate borders of *3a* hotspot, ticks on the *x*-axis represent polymorphisms between Col and Ws. Black arrows represent genes, dashed horizontal line - male chromosome 3 average crossover frequency. *3a-P* target region is shaded in blue and positions of individual guide RNAs are shown as black ticks. **(D)** As in **(C)** but in the presence of *gRNA-P* gRNAs. **(E)** Gene conversion events detected in Col/Ws *MTOPVIB-dCas9 gRNA-P* F_2_ population. DNA sequence polymorphisms (SNPs and InDels) are shown as black ticks at the top of the plot. Maximum and minimum gene conversion tracts shown as black lines and blue rectangles, respectively.

## Discussion

In this study we targeted Arabidopsis MTOPVIB, which is essential for initiation of meiotic DSBs, to the crossover hotspot *3a* via CRISPR. We confirmed that the MTOPVIB-dCas9 translational fusion functionally complements the *mtopvib* mutant. We also confirmed the functionally of the gRNAs we used via catalytically active Cas9 mutagenesis at target loci. The *3a* crossover hotspot is a 5.8 kb sub-telomeric interval with recombination up to ~20 times higher than the chromosome average (Yelina et al. 2012, 2015; Choi et al. 2013). We chose *3a* hotspot as a target firstly because *3a* crossover rates are amenable to manipulation. For example, we have previously shown that recruitment of heterochromatic features, including DNA methylation and H3K9me2, reduces *3a* crossover rates by ~2-3 fold (Yelina et al. 2015). *3a* crossover rates are also not at their maximum level in wild type, as genome-wide loss of DNA methylation results in a ~40% increase in *3a* crossover frequency (Yelina et al. 2012). Secondly, because studies in budding yeast have shown that tethering SPO11 to recombination hot spots leads to a more robust DSB induction compared to targeting SPO11 to DSB cold spots (Sarno et al. 2017; Ito et al. 2014). Mapping of SPO11-1-oligonucleotides in Arabidopsis has revealed that they accumulate at higher levels in nucleosome-free regions (Choi et al. 2018). Therefore, we chose to target MTOPVIB-dCas9 to three locations within *3a* that vary in the nucleosome occupancy levels. We observe very modest and statistically insignificant increases to crossover frequencies and a very similar crossover topology within *3a* in the presence of *gRNA-P*, *gRNA-B* or *gRNA-I* compared to the negative controls.

To explain these results, it is important to note that although DSBs and crossovers correlate positively at the chromosome-scale, there are also regions where the relationship is less strong (He et al. 2017; Choi et al. 2018). Meiotic DSB repair in Arabidopsis is a multi-step process with only ~5-10% of DSBs typically maturing into crossovers (Choi et al. 2018; Serrentino and Borde 2012; Copenhaver et al. 1998; Chelysheva et al. 2012, 2007; Giraut et al. 2011; Salomé et al. 2012). This is in stark contrast to budding yeast where over a half of meiotic DSBs are repaired as crossovers (Pan et al. 2011; Mancera et al. 2008). This could explain why tethering of SPO11 to DSB hotpots in yeast robustly increases recombination (Sarno et al. 2017). Another explanation for our results is that although MTOPVIB is targeted to the *3a* locus, any additional DSBs are repaired via non-crossover pathways. Counter to this, we also did not measure a significant increase in gene conversions in MTOPVIB-Cas9. Specifically, we observed two gene conversion events at a frequency of ~0.21% per SNP each following targeting of MTOPVIB-dCas9 by *gRNA-B* only. Both gene conversion events occurred 1.3-3 kb downstream of the *gRNA-B* target site and did not overlap with each other or *gRNA-B*. The gene conversion frequency we observed is similar to the previously reported Arabidopsis gene conversion frequencies of 0.017-0.55% per SNP at a meiotic crossover hotspot (Drouaud et al. 2013). However, it is important to note that non-crossovers are only detectable when they lead to gene conversions. In Arabidopsis, detectable gene conversion rates are extremely low, with an average of 1.7 per meiosis and are around 100-150 base pairs in length (Lu et al. 2012; Wijnker et al. 2013). The *3a* SNPs measured for gene conversion are spaced 0.7 to 2 kilobase apart. Hence, it is possible that many gene conversions that occur within these intervals would not be detectable. Alternatively, the lack of increased gene conversion frequency upon targeting MTOPVIB-dCas9 may imply that meiotic DSB repair occurs using the sister chromatid as a template (Cifuentes et al. 2013; Yao et al. 2020).

In conclusion, we show that targeting MTOPVIB to an Arabidopsis meiotic recombination hotspot via CRISPR/dCas9 leads to no significant changes in *3a* crossover frequency or pattern. This highlights the complexity of plant meiotic recombination control. We propose that combined recruitment of crossover designation factors and modulation of DSB repair pathways to favour crossovers could be alternative strategies to boost plant crossovers in specific genome locations.

## Materials and methods

### Plant material and genotyping

Arabidopsis lines used in this study were Col-0, *mtopvib-1* (EDA42 line, Ws-4 accession), *mtopvib-2* (GABI_314G09, Col-0 accession) (Vrielynck et al. 2016), *CTL 2.10* and *CTL 5.1* (Wu et al. 2015), which were obtained from the Eurasian Arabidopsis Stock Centre (uNASC) and Arabidopsis Biological Resource Centre (ABRC). Plants were grown under long day conditions (16 h light/8 h dark) at 20 °C, as previously described (Yelina et al. 2015). Plant transformation was performed by floral dipping (Zhang et al. 2006). PCR genotyping of *mtopvib-1* and *mtopvib-2* was performed as described (Vrielynck et al. 2016). PCR genotyping of *mtopvib-2* complemented with *MTOPVIB-dCas9* transgenes was performed with MTOP-genot-compl-F and MTOP-genot-compl-R oligonucleotides. Oligonucleotides are listed in Supplemental Table 13.

### *In silico* gRNA design and *in vitro* testing

gRNAs were *in silico* designed using E-CRISP (Heigwer et al. 2014) (http://www.e-crisp.org/E-CRISP), CRISPR-P (Lei et al. 2014) (http://crispr.hzau.edu.cn/CRISPR2) and CRISPR-MIT (crispr.mit.edu, now obsolete) online tools. gRNAs spacer sequences and Arabidopsis genome target coordinates are listed in Supplemental Table 5. gRNA efficiencies of *in silico* designed gRNAs were tested in an *in vitro* CRISPR/Cas9 assay. Briefly, DNA fragments corresponding to *3a-P*, *3a-B* and *3a-I* and harbouring gRNA target sites were PCR-amplified using Arabidopsis genomic DNA and oligonucleotides listed in Supplemental Table 13. gRNAs were obtained by *in vitro* transcription using MEGAscript T7 Transcription Kit (ThermoFisher Scientific). DNA templates for *in vitro* transcription were PCR-amplified using pEn-Chimera vector and oligonucleotides listed in Supplemental Table 13. 300 ng of gRNA transcript was bound to a purified Cas9 protein (New England Biolabs) for 10 minutes at 25° C, followed by the addition of 300 ng of target DNA and incubation at 37 °C for 1 hour. gRNA transcripts were then cleaved by 0.3 μg/μl RNase A for 5 minutes at 37 °C. DNA fragments were separated on a 1.5% agarose gel stained with Midori Green Advance DNA Stain (Geneflow) to visualize the presence or absence of CRISPR/Cas9-induced target DNA cleavage. gRNAs that led to target DNA cleavage in *in vitro* assays were used to generate constructs for Arabidopsis transformation.

### Cloning

To generate *MTOPVIB-dCas9*, a full genomic sequence of *MTOPVIB* (At1g60460) including a 2385 bp region upstream of the ATG start codon and a 294 bp region downstream of the TAG stop codon was PCR amplified with oligonucleotides MTOPVI-Prom-SalI-F and MTOPVI-Term-NotI-R and cloned between *Sal*I and *Not*I restriction endonuclease sites into pGreen0029 vector (Addgene), to yield the pGreen-gMTOPVIB construct. A *Xba*I restriction endonuclease site in the 7^th^ intron of *MTOPVIB* was mutagenized by digesting pGreen-gMTOPVIB with *Xba*I restriction enzyme, end-filling the resulting 5’ overhang using Klenow fragment and religating to yield pGreen-gMTOPVIBΔXbaI. An *Asc*I restriction site was introduced in front of the ATG start codon by amplifying a part of the MTOPVIB promoter region with MTOPVI-*Nhe*I-F and MTOPVI-*Asc*I-R oligonucleotides and cloning the resulting fragment into *Nhe*I- and *Nco*I-digested pGreen-AscI-gMTOPVIBΔXbaI. A GGSGGS linker, a nuclear localisation signal, two hemagglutinin (2ˣHA) epitope tags and *Xba*I and *Bam*HI restriction sites were introduced at the C-terminus of *MTOPVIB* upstream of the TAG stop codon by cloning a double-strand DNA fragment resulting from annealing MTOPVIB-C-HA-top and MTOPVIB-C-HA-bottom oligonucleotides into a *Pst*I-digested pGreen-AscI-gMTOPVIBΔXbaI. The resulting construct was called pGreen-gMTOPVIB-C-NLS-2ˣHA.

Catalytically inactive Cas9 (dCas9) was generated via PCR-site-directed mutagenesis. Briefly, Cas9 coding sequence was amplified from hSpCas9 plasmid, kindly provided by Prof Jian-Kang Zhu (Feng et al. 2013), in a multiplex PCR reaction using Cas9-1stMut-F, dCas9-1stMut-R, dCas9-2ndMut-F, dCas9-2ndMut-R primers and a Phusion DNA polymerase. Following PCR amplification methylated template plasmid DNA carrying wild type Cas9 was digested with *Dpn*I restriction endonuclease, PCR products carrying mutated dCas9 were ligated and transformed into *E.coli* DH5α strain. Mutations leading to D10A and H840A amino acid substitutions in the Cas9 coding sequence were confirmed by Sanger sequencing. Next, dCas9 was PCR amplified with dCas9-XbaI-F and dCas9-BamHI-R oligonucleotides and cloned into *Xba*I- and *Bam*HI-digested pGreen-gMTOPVIB-C-NLS-2HA to yield MTOPVIB-dCas9. Oligonucleotide sequences are provided in Supplemental Table 13.

To generate Cas9-gRNA-P, Cas9-gRNA-B, Cas9-gRNA-I and Cas9-non-3a-gRNA constructs, 6ˣ(pre-tRNA-gRNA) PCR products were amplified using oligonucleotides listed in the Supplemental table 13 and as described (Xie et al. 2015), digested with *Fok*I restriction endonuclease and cloned into *Bbs*I-digested pEn-Chimera vector (kindly provided by Prof Holger Puchta) behind the Arabidopsis U6 (AtU6) promoter. Fragments containing AtU6:6ˣ(pre-tRNA-gRNA) were transferred from pEn-Chimera into binary pDe-CAS9 vector (kindly provided by Prof Holger Puchta) as described (Schiml et al. 2016).

To generate gRNA-P, gRNA-B, gRNA-I and non-3a-gRNA constructs, 6ˣ(pre-tRNA-gRNA) fragments were PCR amplified as described above, digested with *Fok*I restriction endonuclease and cloned into *Bbs*I-digested pChimera vector, kindly provided by Prof Holger Puchta, behind *AtU6* promoter. Fragments containing AtU6:6ˣ(pre-tRNA-gRNA) were excised from the resulting vectors with *Avr*II restriction endonuclease and cloned into a *Xba*I-digested binary vector pGreen0229.

### Detection of CRISPR/Cas9-induced mutations

*3a-P*, *3a-B* and *3a-I* genetic intervals were PCR amplified from Arabidopsis T_1_ genomic DNA or wild type Col using oligonucleotides listed in Supplemental Table 13. The resulting PCR products were separated on a 1% agarose gel and stained with Midori Green Advance DNA Stain (Geneflow) to visualize full-length and deletion products. The latter were excised and extracted from an agarose gel and subject to Sanger sequencing. Deletion products that could not be resolved by agarose gels were cloned into pGem-T-easy vector (Promega) following the manufacturer’s protocol and individual clones were subject to Sanger sequencing. CRISPR/Cas9-induced mutations in *CLE10*, *CLV3* and *GL1* destroyed *Bsu*36I, *Bsp*HI and *Dde*I restriction endonuclease sites, respectively. To detect CRISPR/Cas9-induced mutations in these genes, DNA fragments harbouring gRNA target sequences were PCR-amplified and digested with the above restriction endonucleases. The resulting products were separated on 1% agarose gels and stained with Midori Green Advance DNA Stain (Geneflow). T7 endonuclease I (New England Biolabs) assays were used to detect CRISPR/Cas9-induced mutations in *CLE9*, *FWA* and *eIF(iso)4E* as described (Pyott et al. 2016).

### Seed fluorescent measurement of crossovers

Seed fluorescent measurements of crossovers in *CTL 2.10* and *CTL 5.1* intervals was performed as described (Yelina et al. 2015), using CellProfiler (Carpenter et al. 2006).

### Pollen typing

Pollen typing for *3a* crossover hotspot was performed as previously described (Yelina et al. 2015).

### RT-PCR detection of gRNA transcripts

RNA was extracted from closed buds of two independent pools of F_1_ individuals used for ‘pollen typing’ using PureZOL™ RNA Isolation Reagent (Bio-Rad) according to the manufacturer’s protocol. 10 μg of total RNA was treated with TURBO DNase (ThermoFisher Scientific) and reverse-transcribed in the presence or absence (negative control) of SuperScript IV enzyme (ThermoFisher) using random hexamer primers, according to the manufacturers’ protocols. A 1:20 dilution of the resulting cDNA was PCR-amplified using oligonucleotides listed in Supplemental Table 13. The resulting products were resolved on a 2% agarose gel stained with Midori Green Advance DNA Stain (Geneflow).

### Kompetitive Allele-Specific PCR (KASP) Assay

Arabidopsis genomic DNA was extracted as described (Edwards et al. 1991). Kompetitive Allele-Specific PCR (KASP) Assay was performed following the manufacturer’s protocol using KASP master mix (Bioresearch Technologies) and oligonucleotides are listed in Supplemental table 13. Reactions were run on a CFX real-time PCR system (Bio-Rad), allele discrimination was performed using the manufacturer’s software.

## Data availability

All plasmids, reagents and Arabidopsis transgenic lines generated in this study are available upon request. Supplemental Tables S1-S13 contain raw data used for fertility and crossover frequency scoring, as well as genomic positions of *3a* crossover hotspot, guide RNAs, SNPs and oligonucleotides used in this study. Supplemental figures S1-S3 contain CRISPR/Cas9 gene editing analysis and supplemental figure S4 contains confirmation of gRNA expression.

## Acknowledgements

Research was supported by a Broodbank Fellowship (N.E.Y.), a Gatsby Grant to Exceptional Researchers (N.E.Y and I.R.H) and a BBSRC-IPA grant BB/N007557/1 with Meiogenix. We thank Professor Holger Puchta for pEn-Chimera, pChimera and pDe-CAS9 vectors, Professor Jian-Kang Zhu for hSpCas9 plasmid, Dr Wei Jiang for advice on the *in vitro* CRISPR/Cas9 and KASP assays, Piotr Wlodzimierz for an automated CellProfiler fluorescent seed scoring pipeline, Prof Alain Nicolas and Daniel Holland for discussions, Mel Steer, Emma Jackson and James Barlow for technical support.

---

**Figure S1.**
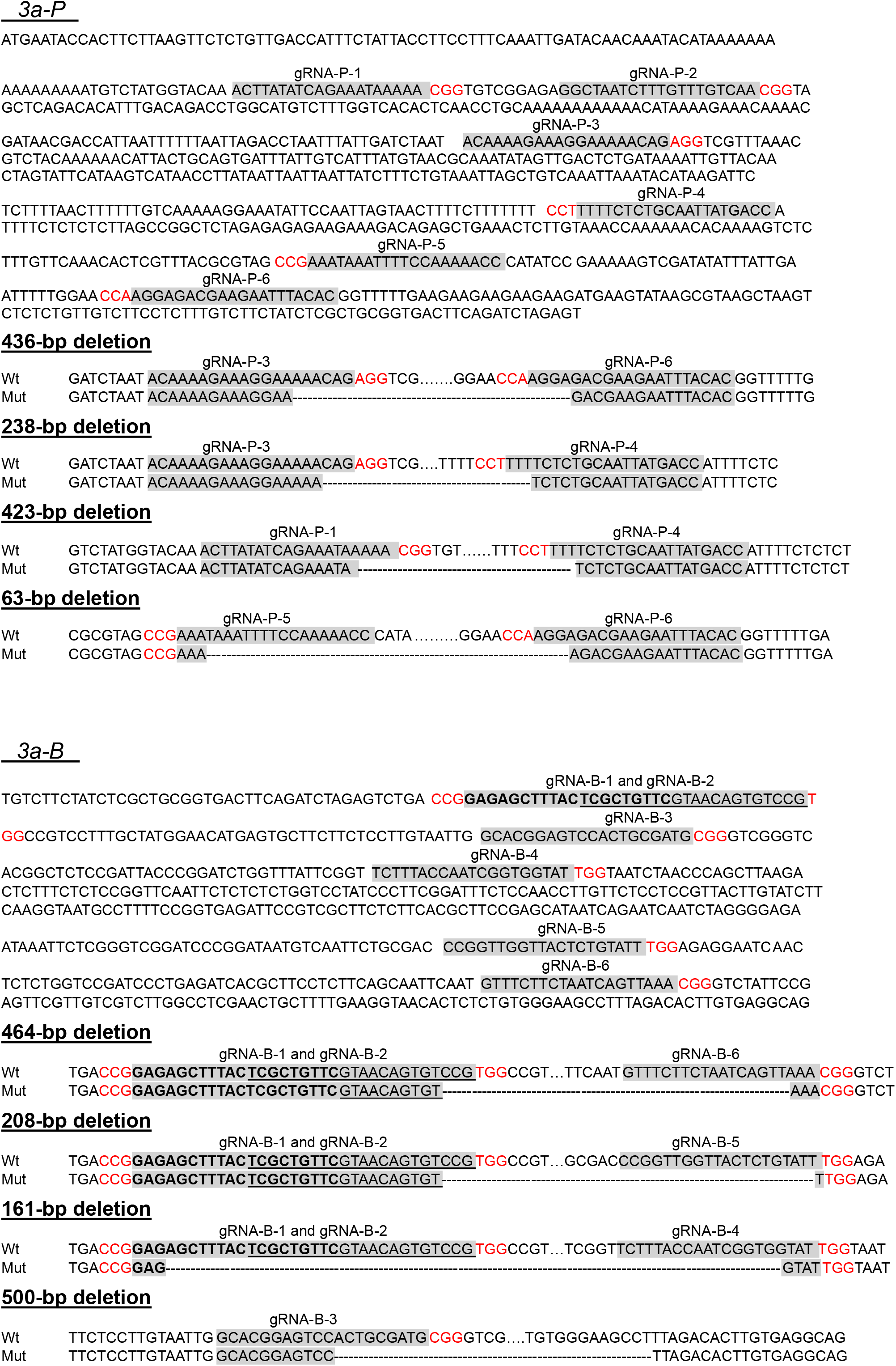
Sanger sequencing analysis of gene editing events in *3a* crossover hotspot. Nucleotide sequences of each of the target regions (*3a-P*, *3a-B* and *3a-I*) are shown, gRNA target sites highlighted in grey and protospacer adjacent motif, PAM, in red. To distinguish between overlapping gRNA-B-1 and gRNA-B-2, in addition to highlighting them in grey, one is shown bold and the other is underlined. Dots indicate wild type genomic sequence not shown due to space constraints, dashes indicate sequences deleted via CRISPR/Cas9-mediated gene editing. In the case where CRISPR/Cas9 editing causes an 8 bp insertion, dashes indicate a corresponding missing sequence in the wild type.

**Figure S2.**
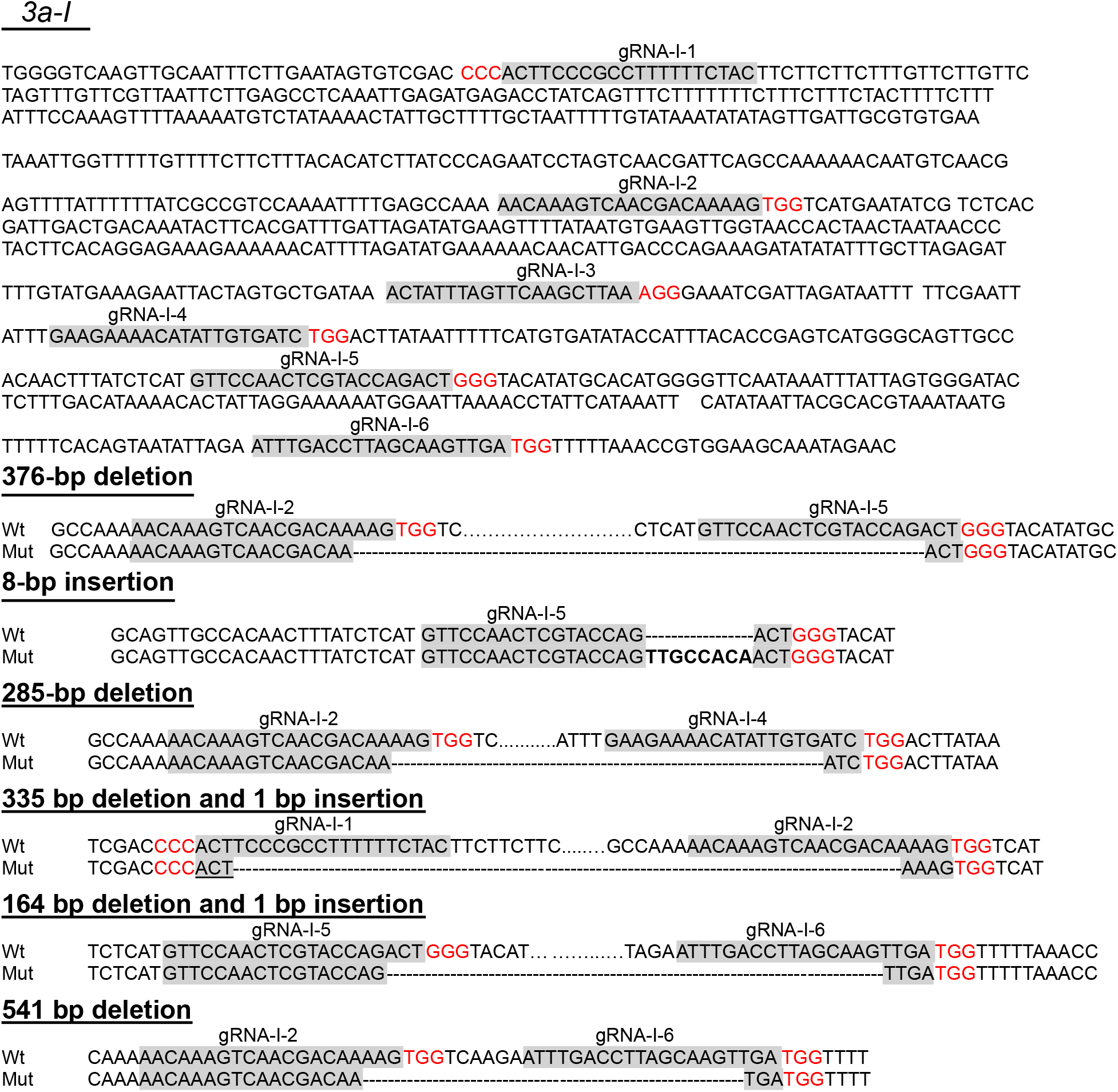
Sanger sequencing analysis of gene editing events in *3a* crossover hotspot. Nucleotide sequences of each of the target regions (*3a-P*, *3a-B* and *3a-I*) are shown, gRNA target sites highlighted in grey and protospacer adjacent motif, PAM, in red. To distinguish between overlapping gRNA-B-1 and gRNA-B-2, in addition to highlighting them in grey, one is shown bold and the other is underlined. Dots indicate wild type genomic sequence not shown due to space constraints, dashes indicate sequences deleted via CRISPR/Cas9-mediated gene editing. In the case where CRISPR/Cas9 editing causes an 8 bp insertion, dashes indicate a corresponding missing sequence in the wild type.

**Figure S3.**
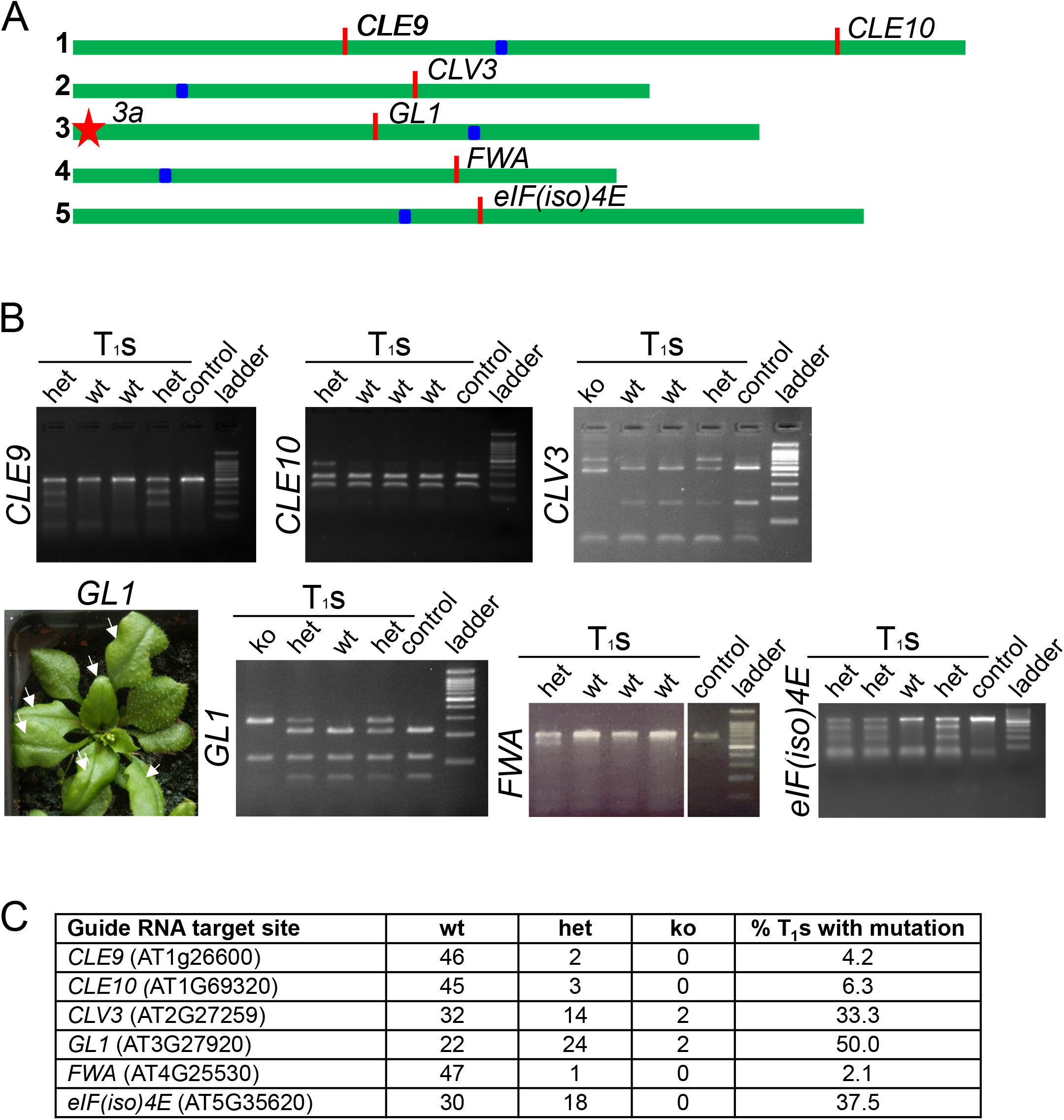
Testing the efficiency of guide RNAs targeting outside *3a* crossover hotspot. **(A)** Schematic representation of the five Arabidopsis chromosomes and gRNA positions. Chromosomes - green bars, blue rectangles - centromeres, red ticks - genes targeted with CRISPR/Cas9, star - 3a hotspot (shown for reference). **(B)** Constructs harbouring guide RNAs targeting six Arabidopsis genes (*CLE9*, *CLE10*, *CLV3*, *GL1*, *FWA* and *eIF(iso)4E*)) and a catalytically active Cas9 were transformed into wild type Col. T_1_ progenies were selected for the presence of *gRNA-Cas9* transgenes and tested for gene editing events. Representative Midori-green-stained agarose gel analyses of CRISPR/Cas9-mediated gene editing in T_1_ leaf tissue. Mutations introduced by CRISPR/Cas9 either destroy a restriction endonuclease recognition site (*Bsu36*I in *CLE10*, *BspH*I in *CLV3* and *Dde*I in *GL1*) resulting in higher molecular weight products compared to wild type control (untransformed Col) or introduce a mismatch that is recognised by T7 Endonuclease I resulting in endonucleolytic cleavage and lower molecular weight product(s) compared to untransformed Col control (in *CLE9*, *eIF(iso)4E* and *FWA*). Different leaf sectors of the analysed T_1_ plants were mosaic for CRISPR/Cas9-mediated gene editing events and had one (‘het’), two (‘ko) or neither (‘wt’) gene edited allele(s). This was visually demonstrated for *GL1* responsible for trichome development. Leaf segments showing absence of trichomes (white arrows) had both *GL1* alleles mutated by CRISRP/Cas9. **(C)** Summary table showing number and percentage of T_1_s affected by CRISPR/Cas9 gene editing.

**Figure S4.**
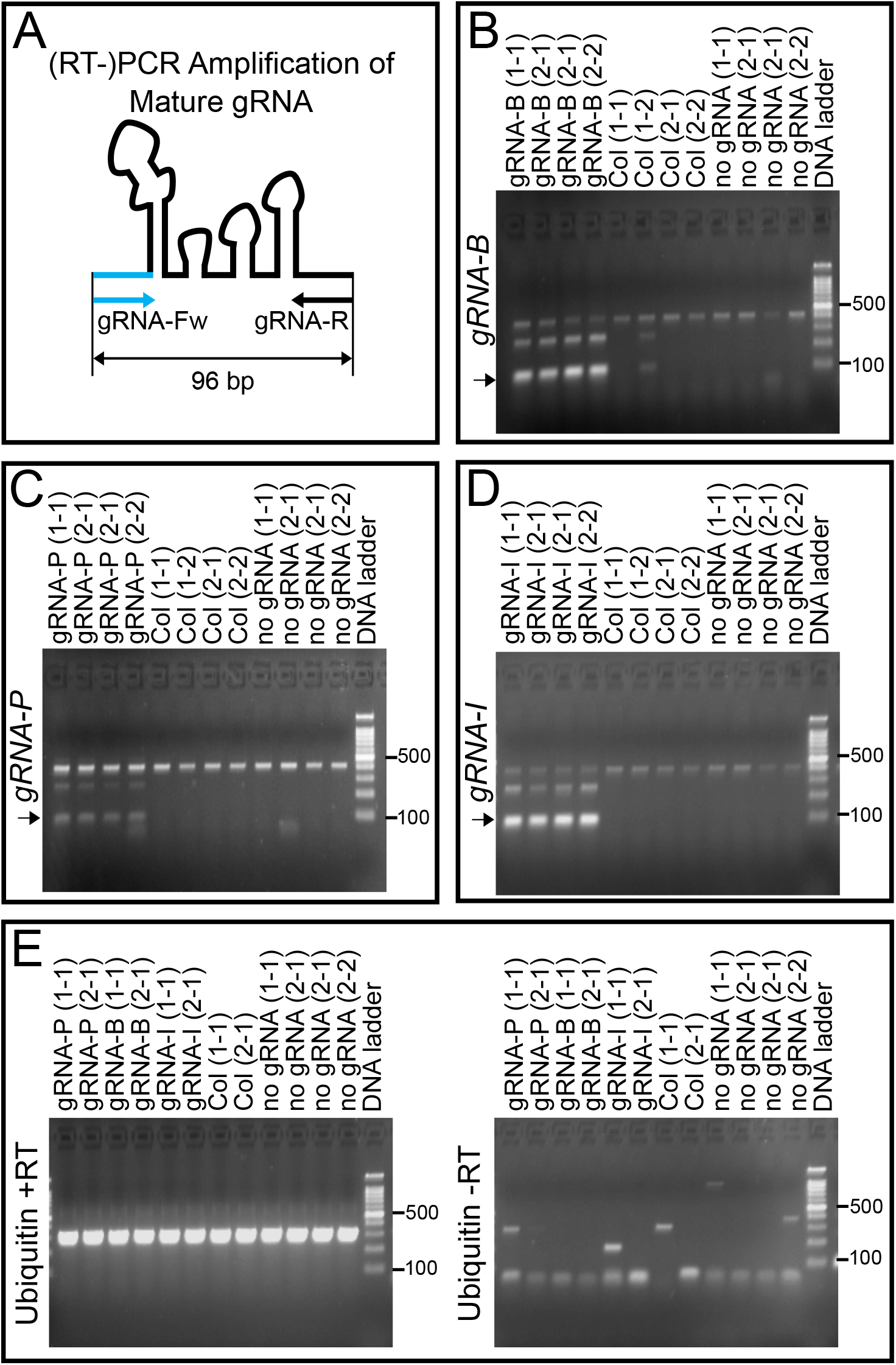
Detection of mature gRNAs via RT-PCR. **(A)** Schematic representation of a mature gRNA. An (RT-)PCR assay to detect mature gRNAs uses a ‘universal’ reverse primer complementary to the 3’ end of the gRNA scaffold (black arrow) and a guide-RNA-spacer-specific forward primer, light blue arrow. **(B)** Detection of one of the six mature gRNAs targeting *3a-B* by RT-PCR in Col/Ws *mtopvib MTOPVIB-dCas9 gRNA-P* F_1_ population. Black arrow points to the mature gRNA-specific PCR product resolved on a Midori-green-strained 2% agarose gel. Col and ‘no gRNA’ are negative controls. Two biological and two technical replicates were done for each genotype. Higher than 96-bp molecular weight bands are non-specific products. **(C)** As in **(B)** but for *gRNA-P*. **(D)** As in **(B)** but for *gRNA-I*. Two to three out of the six multiplexed gRNAs for each of the target regions was tested, but data for one gRNA is shown. **(E)** Ubiquitin was used as a ‘+RT’ and ‘−RT’ controls.

## SUPPLEMENTAL TABLES

**Table 1.** Seed sets in Col, *mtopvib* and complementing *MTOPVIB-dCas9 mtopvib* lines. Two-tailed t-test was used to calculate the *P* values

| Genotype | Total siliques scored | Total seeds scored | Average | Standard Deviation | *P* value |
| --- | --- | --- | --- | --- | --- |
| Col | 10 | 645 | 64.5 | 8.2 |  |
| *mtopvib* | 56 | 105 | 1.9 | 1.7 | <0.00001 |
| *MTOPVIB-dCas9 mtopvib* | 7 | 463 | 66.1 | 3.3 | 0.63 |

**Table 2.** Genetic distances of *CTL2.10* in wild type and *MTOPVIB-dCas9 mtopvib*. cM were calculated using the formula: cM = 100 × (1 – [1-2(*N_G_*+*N_R_*)/*N_T_*] ^½^), where *N_G_* is a number of green-alone fluorescent seeds, *N_R_* is a number of red-alone fluorescent seed and *N_T_* is the total number of seeds counted.

| Genotype | Green-alone | Red-alone | Both red and green | Neither red or green | Total | CTL 2.10, cM |
| --- | --- | --- | --- | --- | --- | --- |
| Wild type | 140 | 152 | 1744 | 467 | 2503 | 12.44 |
| Wild type | 133 | 129 | 1611 | 453 | 2326 | 11.98 |
| Wild type | 131 | 147 | 1649 | 447 | 2374 | 12.49 |
| Wild type | 131 | 166 | 1727 | 440 | 2464 | 12.88 |
| Wild type | 134 | 161 | 1734 | 477 | 2506 | 12.56 |
| Wild type | 145 | 163 | 1621 | 437 | 2366 | 14.00 |
| Wild type | 109 | 169 | 1695 | 449 | 2422 | 12.23 |
| Wild type | 136 | 139 | 1648 | 445 | 2368 | 12.38 |
| *MTOPVIB-dCas9 mtopvib* | 120 | 153 | 1665 | 449 | 2387 | 12.18 |
| *MTOPVIB-dCas9 mtopvib* | 138 | 174 | 1678 | 423 | 2413 | 13.90 |
| *MTOPVIB-dCas9 mtopvib* | 124 | 143 | 1597 | 413 | 2277 | 12.51 |
| *MTOPVIB-dCas9 mtopvib* | 131 | 146 | 1588 | 416 | 2281 | 12.99 |
| *MTOPVIB-dCas9 mtopvib* | 127 | 157 | 1681 | 463 | 2428 | 12.47 |
| *MTOPVIB-dCas9 mtopvib* | 145 | 169 | 1675 | 405 | 2394 | 14.11 |
| *MTOPVIB-dCas9 mtopvib* | 148 | 159 | 1648 | 431 | 2386 | 13.82 |
| *MTOPVIB-dCas9 mtopvib* | 147 | 174 | 1656 | 425 | 2402 | 14.40 |
| *MTOPVIB-dCas9 mtopvib* | 112 | 143 | 1627 | 417 | 2299 | 11.79 |
| *MTOPVIB-dCas9 mtopvib* | 133 | 164 | 1656 | 493 | 2446 | 12.99 |
| *MTOPVIB-dCas9 mtopvib* | 127 | 141 | 1599 | 433 | 2300 | 12.42 |

**Table 3.** Genetic distances of *CTL5.1* in wild type and *MTOPVIB-dCas9 mtopvib*. cM were calculated using the formula: cM = 100 × (1 – [1-2(*N_G_*+*N_R_*)/*N_T_*] ^½^), where *N_G_* is a number of green-alone fluorescent seeds, *N_R_* is a number of red-alone fluorescent seed and *N_T_* is the total number of seeds counted.

| **Genotype** | **Green-alone** | **Red-alone** | **Both red and green** | **Neither red or green** | **Total** | **CTL 5.1, cM** |
| --- | --- | --- | --- | --- | --- | --- |
| Wild type | 220 | 233 | 1455 | 353 | 2261 | 22.59 |
| Wild type | 276 | 264 | 1477 | 364 | 2381 | 26.08 |
| Wild type | 242 | 246 | 1532 | 332 | 2352 | 23.51 |
| Wild type | 264 | 275 | 1586 | 356 | 2481 | 24.80 |
| Wild type | 250 | 251 | 1661 | 379 | 2541 | 22.18 |
| Wild type | 280 | 237 | 1633 | 343 | 2493 | 23.50 |
| Wild type | 239 | 257 | 1648 | 355 | 2499 | 22.34 |
| Wild type | 193 | 225 | 1217 | 254 | 1889 | 25.34 |
| Wild type | 264 | 241 | 1559 | 363 | 2427 | 23.59 |
| *MTOPVIB-dCas9*<br>*mtopvib* | 229 | 234 | 1479 | 334 | 2276 | 22.98 |
| *MTOPVIB-dCas9*<br>*mtopvib* | 271 | 248 | 1562 | 306 | 2387 | 24.82 |
| *MTOPVIB-dCas9*<br>*mtopvib* | 273 | 272 | 1544 | 349 | 2438 | 25.64 |
| *MTOPVIB-dCas9*<br>*mtopvib* | 251 | 204 | 1354 | 339 | 2148 | 24.08 |
| *MTOPVIB-dCas9*<br>*mtopvib* | 269 | 247 | 1551 | 376 | 2443 | 24.00 |
| *MTOPVIB-dCas9*<br>*mtopvib* | 249 | 212 | 1489 | 327 | 2277 | 22.86 |
| *MTOPVIB-dCas9*<br>*mtopvib* | 259 | 241 | 1515 | 382 | 2397 | 23.66 |
| *MTOPVIB-dCas9*<br>*mtopvib* | 280 | 251 | 1621 | 360 | 2512 | 24.02 |
| *MTOPVIB-dCas9*<br>*mtopvib* | 286 | 228 | 1482 | 343 | 2339 | 25.13 |
| *MTOPVIB-dCas9*<br>*mtopvib* | 288 | 250 | 1542 | 364 | 2444 | 25.18 |
| *MTOPVIB-dCas9*<br>*mtopvib* | 250 | 209 | 1555 | 345 | 2359 | 21.84 |
| *MTOPVIB-dCas9*<br>*mtopvib* | 319 | 240 | 1566 | 388 | 2513 | 25.49 |
| *MTOPVIB-dCas9*<br>*mtopvib* | 254 | 221 | 1639 | 358 | 2472 | 21.53 |
| *MTOPVIB-dCas9*<br>*mtopvib* | 230 | 219 | 1457 | 334 | 2240 | 22.60 |

**Table 4.** Coordinates of *3a* crossover hotspot and genes within *3a*.

| Locus | Chromosome | Start coordinate | End coordinate |
| --- | --- | --- | --- |
| 3a crossover hotspot | 3 | 634,109 | 639,934 |
| AT3G02880 | 3 | 634,465 | 637,284 |
| AT3G02885 | 3 | 637,921 | 639,178 |

**Table 5.**
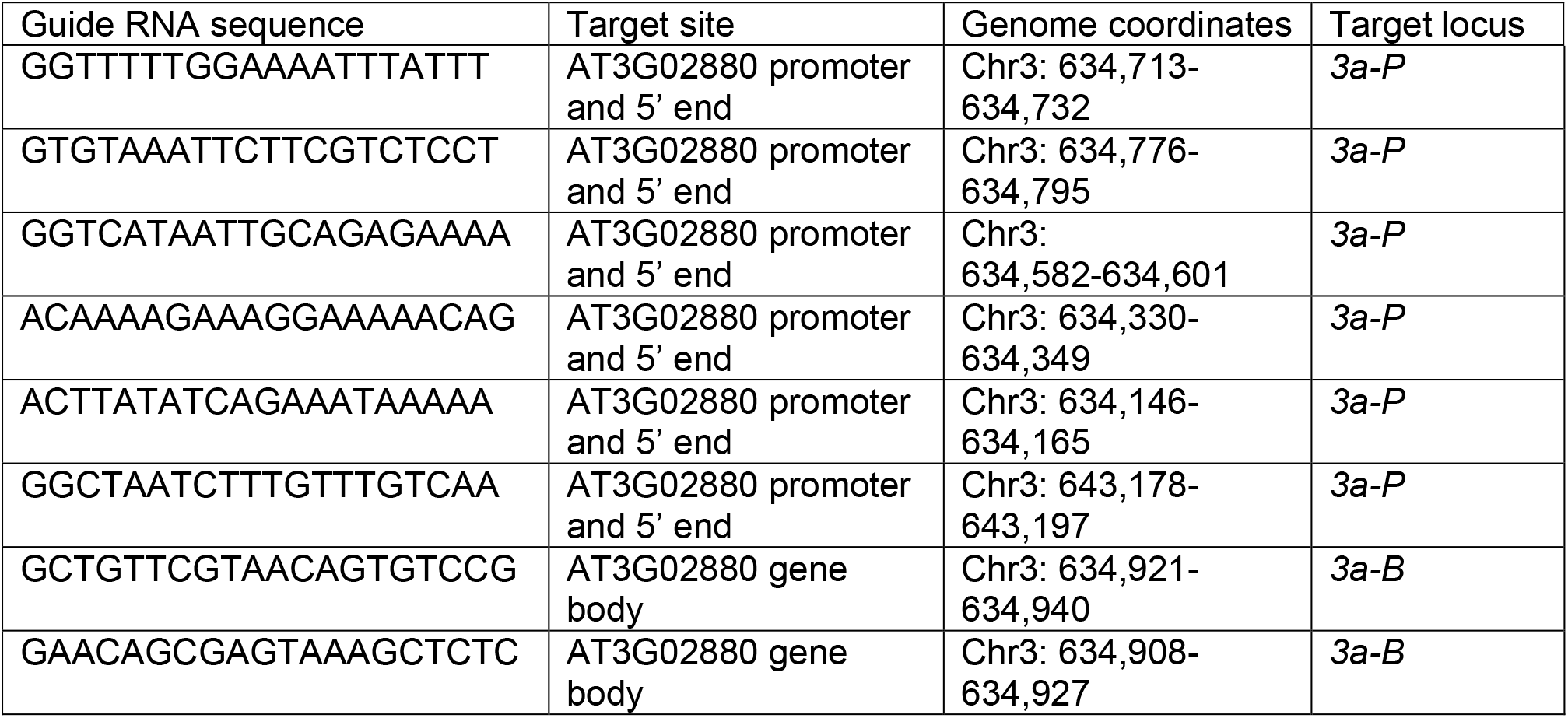
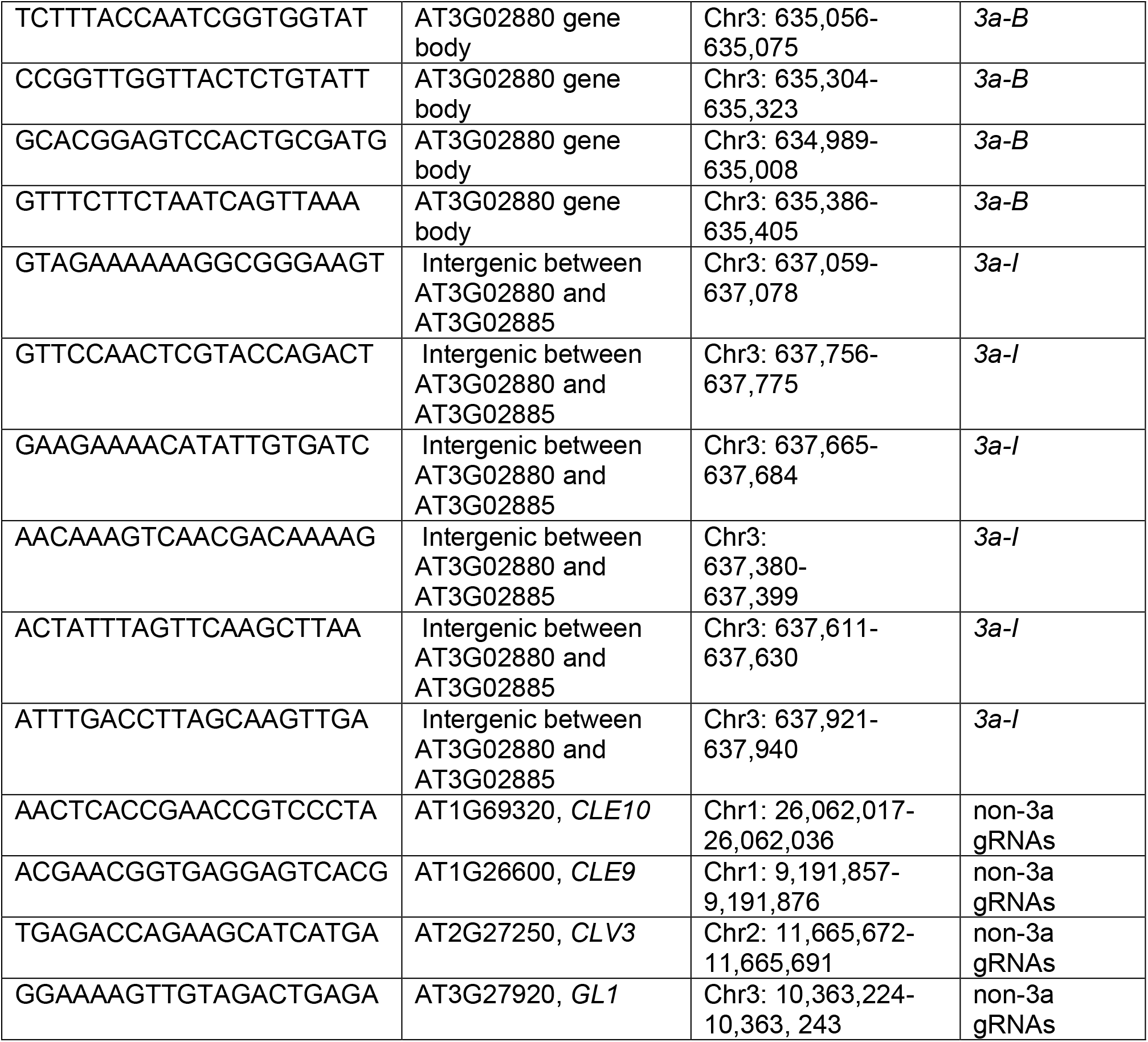
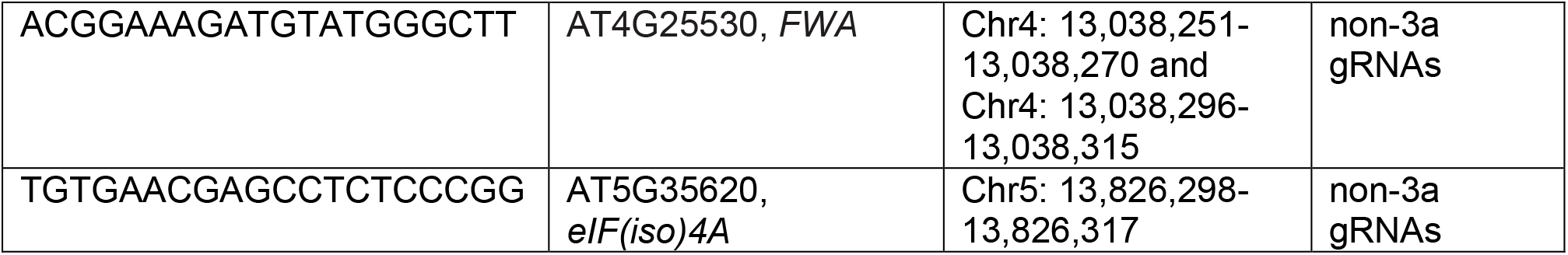
Sequences and coordinates of guide RNAs used in this study.

**Table 6.**
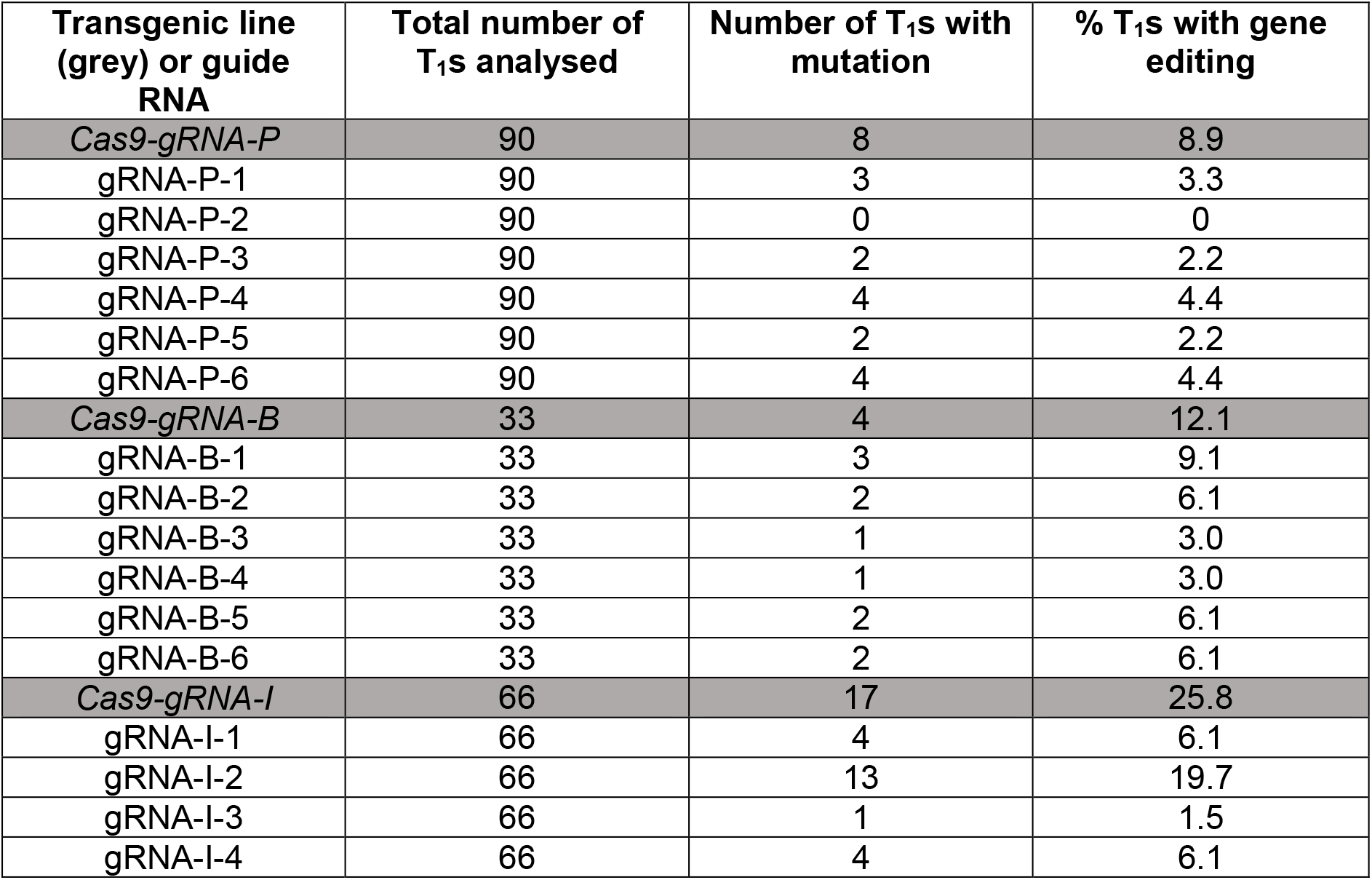
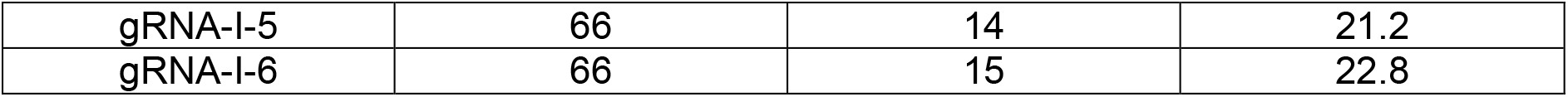
CRISPR/Cas9 gene editing efficiencies in *3a*. Number and percentage of *Cas9-gRNA-P*, *Cas9-gRNA-B* and *Cas9-gRNA-I* T_1_ progeny with gene editing events and gene editing efficiencies for individual guide RNAs.

**Table 7.** CRISPR/Cas9 gene editing efficiencies outside of *3a* hotspot.

| Guide RNA target site | Wild type | One allele affected | Both alleles affected | % T<sub>1</sub>s with mutation |
| --- | --- | --- | --- | --- |
| *CLE9* | 46 | 2 | 0 | 4.2 |
| *CLE10* | 45 | 3 | 0 | 6.3 |
| *CLV3* | 32 | 14 | 2 | 33.3 |
| *GL1* | 22 | 24 | 2 | 50.0 |
| *FWA* | 47 | 1 | 0 | 2.1 |
| *eIF(iso)4E* | 30 | 18 | 0 | 37.5 |

**Table 8.** *3a* crossover frequencies in Col/Ws *MTOPVIB-dCas9 mtopvib* F_1_ populations in the presence or absence of guide RNAs targeting *3a*. Chi-square test was used to calculate *P*-values.

| Genotype | Crossovers per microliter | Parentals per microliter | 3a | Standard Deviation | 3a | *P*-value |
| --- | --- | --- | --- | --- | --- | --- |
| non-3a gRNA | 26.9 | 17412 | 0.155 cM | 0.014 | 26.4 cM/Mb |  |
| no gRNA | 32.5 | 24767 | 0.131 cM | 0.011 cM | 22.3 cM/Mb | 0.56 |
| *gRNA-P* | 45.6 | 30059 | 0.152 cM | 0.014 cM | 25.9 cM/Mb | 0.96 |
| *gRNA-B* | 42.3 | 22375 | 0.189 cM | 0.021 cM | 32.2 cM/Mb | 0.44 |
| *gRNA-I* | 41.2 | 23522 | 0.175 cM | 0.020 cM | 29.8 cM/Mb | 0.64 |

**Table 9.** Fine-scale mapping of *3a* crossover frequencies in Col/Ws *MTOPVIB-dCas9 mtopvib* F_1_ populations in the presence or absence of guide RNAs targeting *3a*.

| Coordinate | Interval Length | CO<br>'no<br>gRNA' | cM/Mb<br>'no<br>gRNA' | CO<br>'non-3a<br>gRNA' | cM/Mb<br>'non-3a<br>gRNA' | CO<br>*gRNA-P* | cM/Mb<br>*gRNA-P* | CO<br>*gRNA-B* | cM/Mb<br>*gRNA-B* | CO<br>*gRNA-I* | cM/Mb<br>*gRNA-I* |
| --- | --- | --- | --- | --- | --- | --- | --- | --- | --- | --- | --- |
| 634109 |  |  |  |  |  |  |  |  |  |  |  |
| 634938 | 829 | 0 | 0.0 | 0 | 0.0 | 0 | 0.0 | 0 | 0.0 | 0 | 0.0 |
| 635457 | 519 | 5 | 15.8 | 2 | 7.8 | 0 | 0.0 | 2 | 8.4 | 2 | 7.7 |
| 635520 | 63 | 0 | 0.0 | 1 | 32.0 | 0 | 0.0 | 1 | 34.5 | 0 | 0.0 |
| 636119 | 599 | 3 | 8.2 | 3 | 10.1 | 2 | 5.6 | 3 | 10.9 | 0 | 0.0 |
| 637283 | 1164 | 6 | 8.4 | 9 | 15.6 | 13 | 18.9 | 14 | 26.1 | 13 | 22.2 |
| 637754 | 471 | 8 | 27.8 | 6 | 25.6 | 8 | 28.7 | 11 | 50.7 | 13 | 54.9 |
| 637900 | 146 | 3 | 33.6 | 0 | 0.0 | 0 | 0.0 | 0 | 0.0 | 3 | 40.9 |
| 637952 | 52 | 0 | 0.0 | 0 | 0.0 | 0 | 0.0 | 0 | 0.0 | 0 | 0.0 |
| 637974 | 22 | 0 | 0.0 | 0 | 0.0 | 0 | 0.0 | 0 | 0.0 | 0 | 0.0 |
| 637988 | 14 | 0 | 0.0 | 0 | 0.0 | 0 | 0.0 | 0 | 0.0 | 0 | 0.0 |
| 638139 | 151 | 4 | 43.4 | 3 | 40.0 | 6 | 67.1 | 3 | 43.2 | 2 | 26.3 |
| 638196 | 57 | 3 | 86.2 | 1 | 35.3 | 1 | 29.6 | 0 | 0.0 | 1 | 34.9 |
| 638483 | 287 | 7 | 39.9 | 8 | 56.1 | 17 | 100.0 | 12 | 90.8 | 20 | 138.6 |
| 638786 | 303 | 14 | 75.7 | 13 | 86.4 | 20 | 111.5 | 10 | 71.7 | 12 | 78.8 |
| 638818 | 32 | 1 | 51.2 | 0 | 0.0 | 1 | 52.8 | 1 | 67.9 | 1 | 62.1 |
| 639370 | 552 | 13 | 38.6 | 25 | 91.2 | 16 | 49.0 | 20 | 78.7 | 12 | 43.2 |
| 639538 | 168 | 13 | 126.7 | 6 | 71.9 | 6 | 60.3 | 10 | 129.3 | 9 | 106.5 |
| 639934 | 462 | 0 | 0.0 | 0 | 0.0 | 0 | 0.0 | 0 | 0.0 | 0 | 0.0 |
| Total | 5825 | 80 | 22.49 | 77 | 26.61 | 90 | 26.09 | 87 | 32.45 | 88 | 30.04 |

**Table 10.**
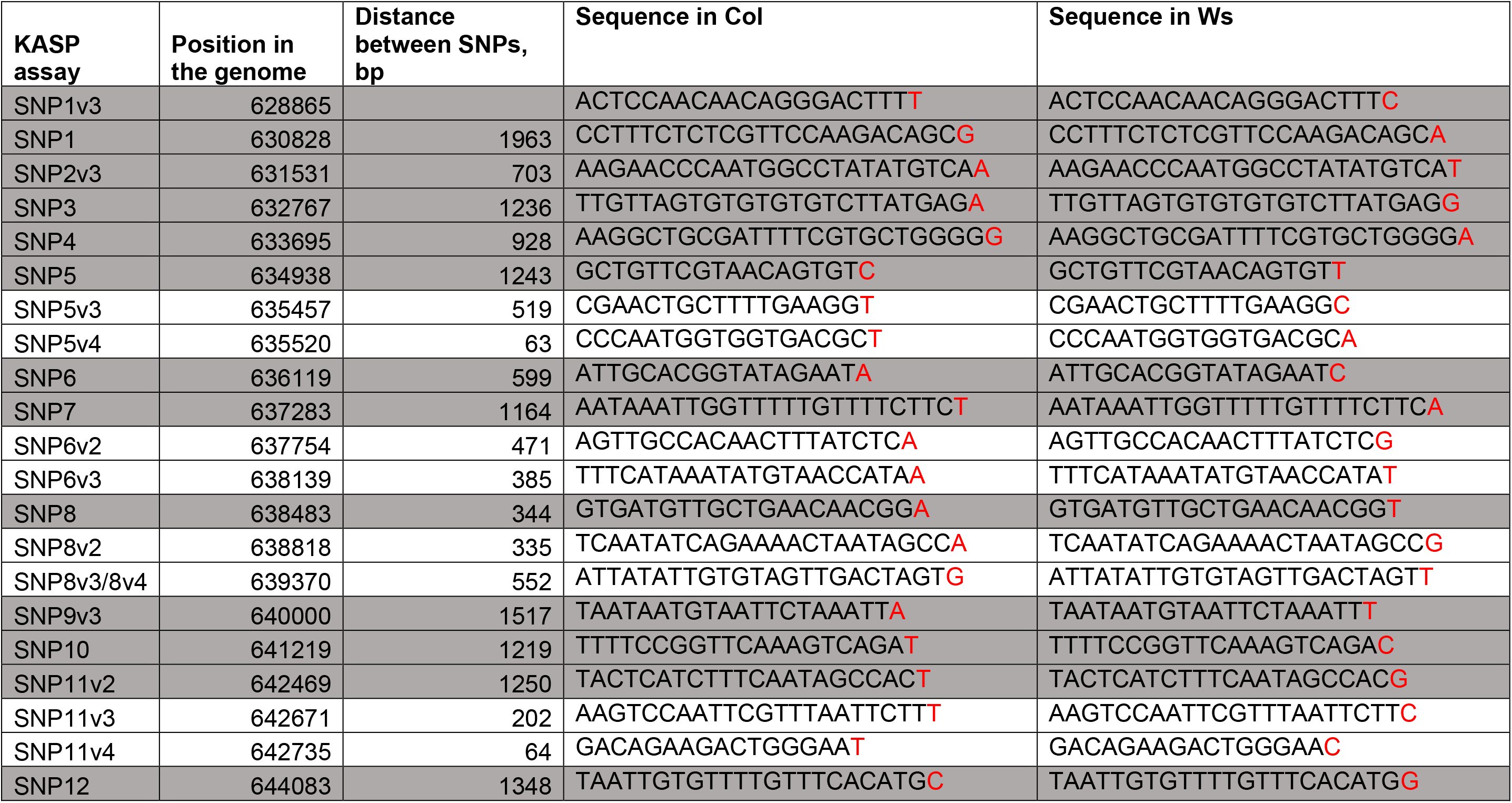
Genomic positions of single nucleotide polymorphisms (SNPs), in red, used for KASP assays. Assays highlighted in grey were used for initial screening of the F_2_ population. Assays in white were used only on individual F_2_ plants to determine gene conversion tract lengths.

**Table 11.**
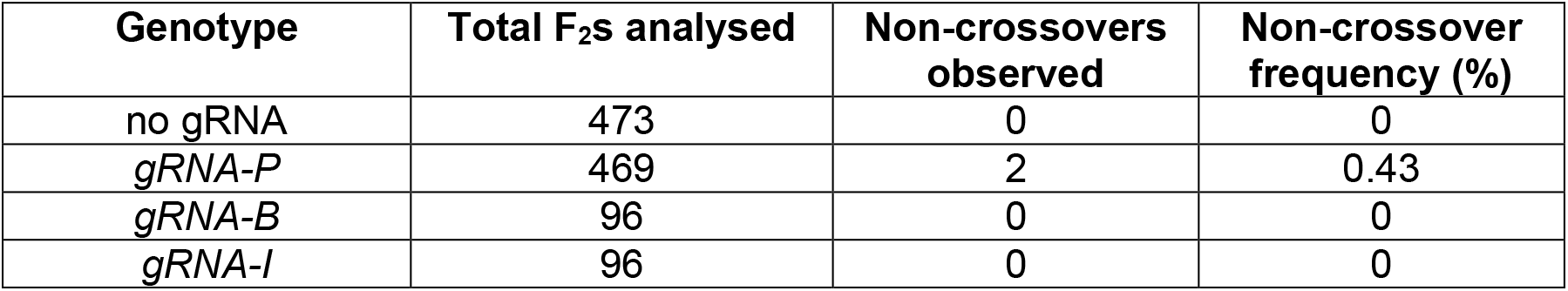
Gene conversion events observed in F_2_ progenies of Col/Ws *MTOPVIB-dCas9 mtopvib* in the presence or absence of guide RNAs targeting *3a*.

**Table 12.** Gene conversion events observed in two individuals (#519 and #343) of F_2_ progeny of Col/Ws *MTOPVIB-dCas9 gRNA-P mtopvib*.

| Polymorphism Coordinate | Interval Length | Genotype |  |
| --- | --- | --- | --- |
|  |  | *gRNA-P*, F<sub>2</sub>#519 | *gRNA-P*, F<sub>2</sub>#343 |
| 634109 |  | Het | Ws |
| 634938 | 829 | Het | Ws |
| 635457 | 519 | Het | Ws |
| 635520 | 63 | Het | Ws |
| 636119 | 599 | Het | Het |
| 637283 | 1164 | Het | Ws |
| 637754 | 471 | Ws | Not tested |
| 637900 | 146 | Ws | Not tested |
| 637952 | 52 | Ws | Not tested |
| 637974 | 22 | Ws | Not tested |
| 637988 | 14 | Ws | Not tested |
| 638139 | 151 | Ws | Not tested |
| 638196 | 57 | Ws | Not tested |
| 638483 | 287 | Ws | Ws |
| 638786 | 303 | Het | Not tested |
| 638818 | 32 | Not tested | Not tested |
| 639370 | 552 | Not tested | Not tested |
| 639538 | 168 | Not tested | Not tested |
| 640000 | 462 | Het | Ws |

**Table 13.**
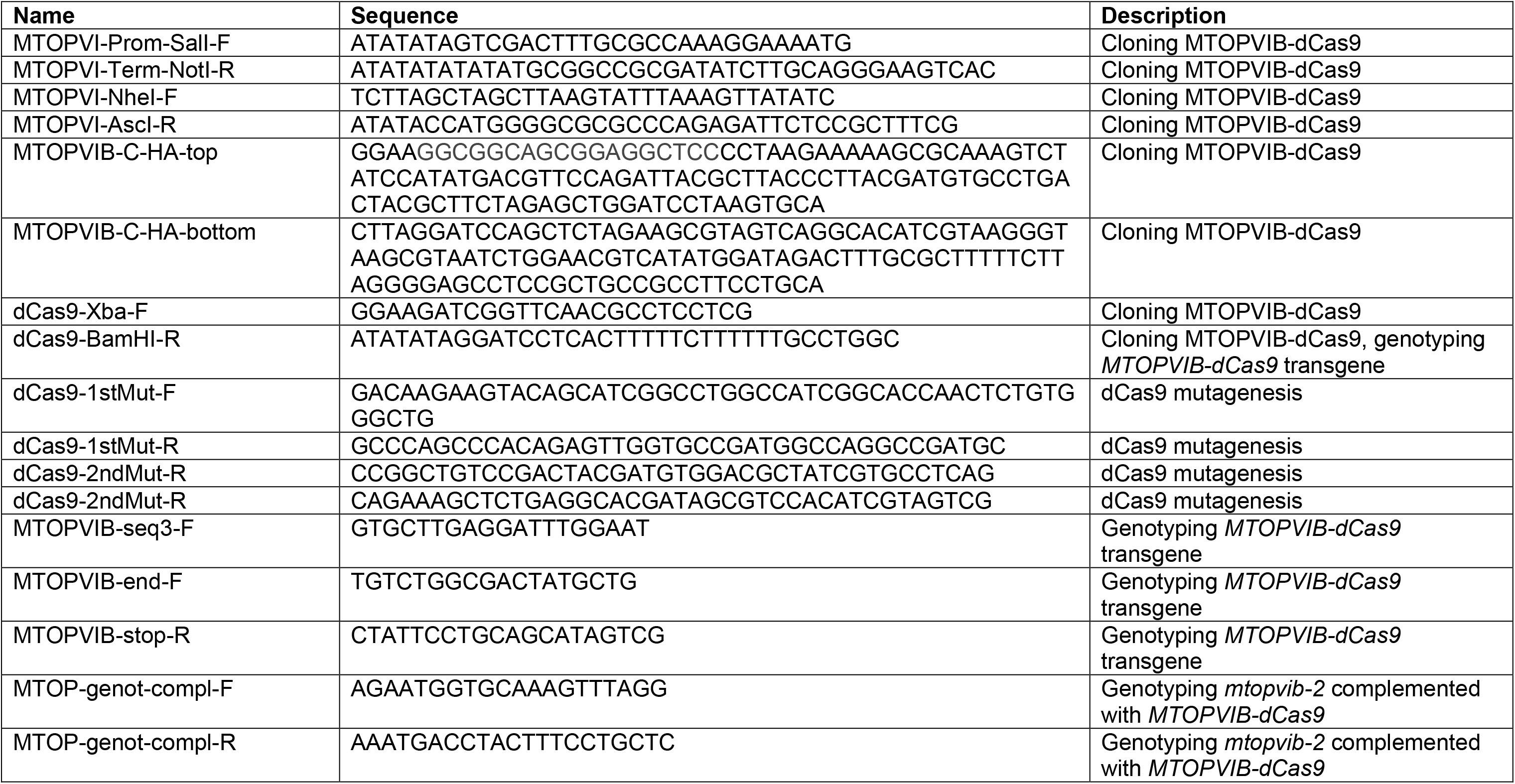
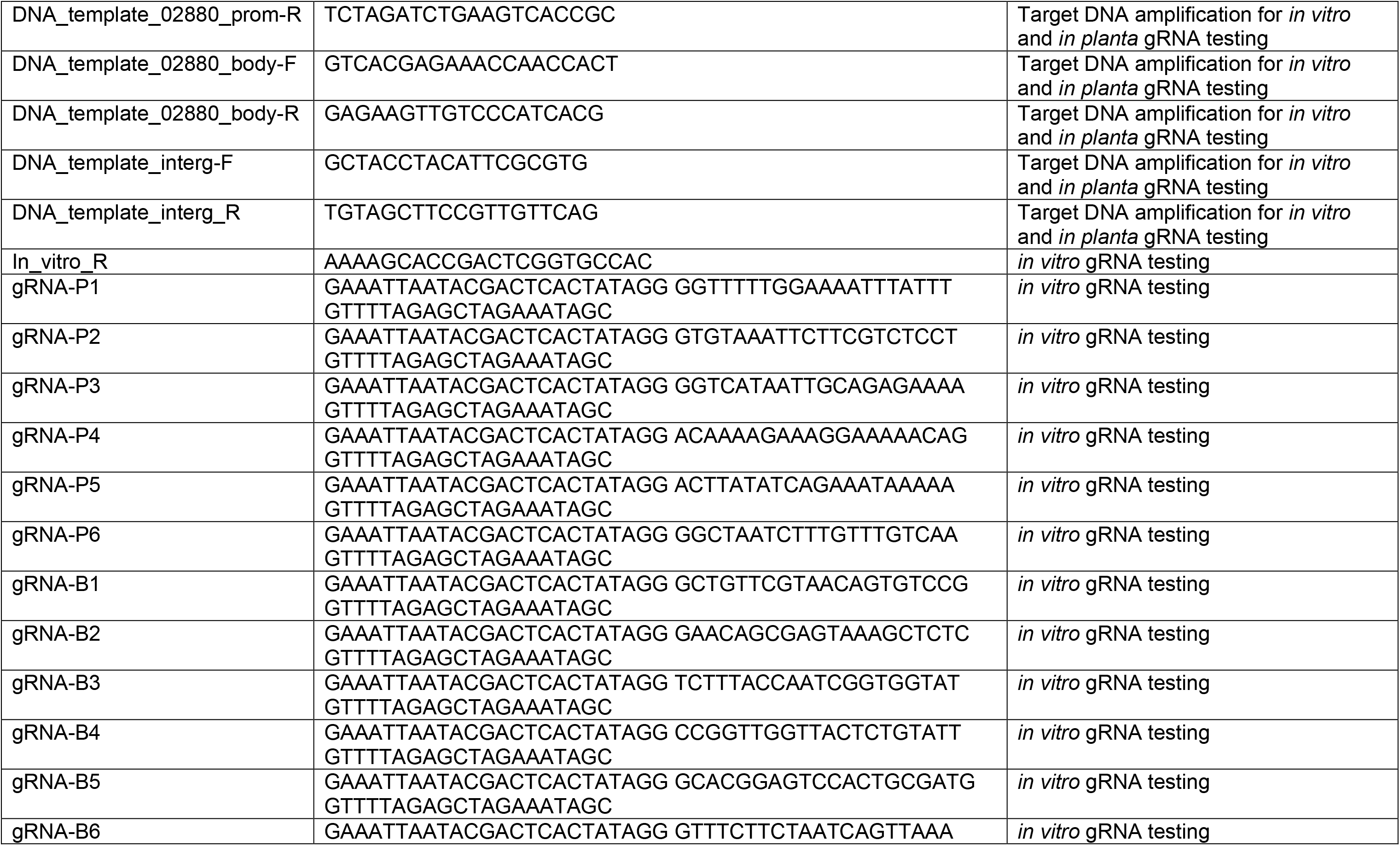
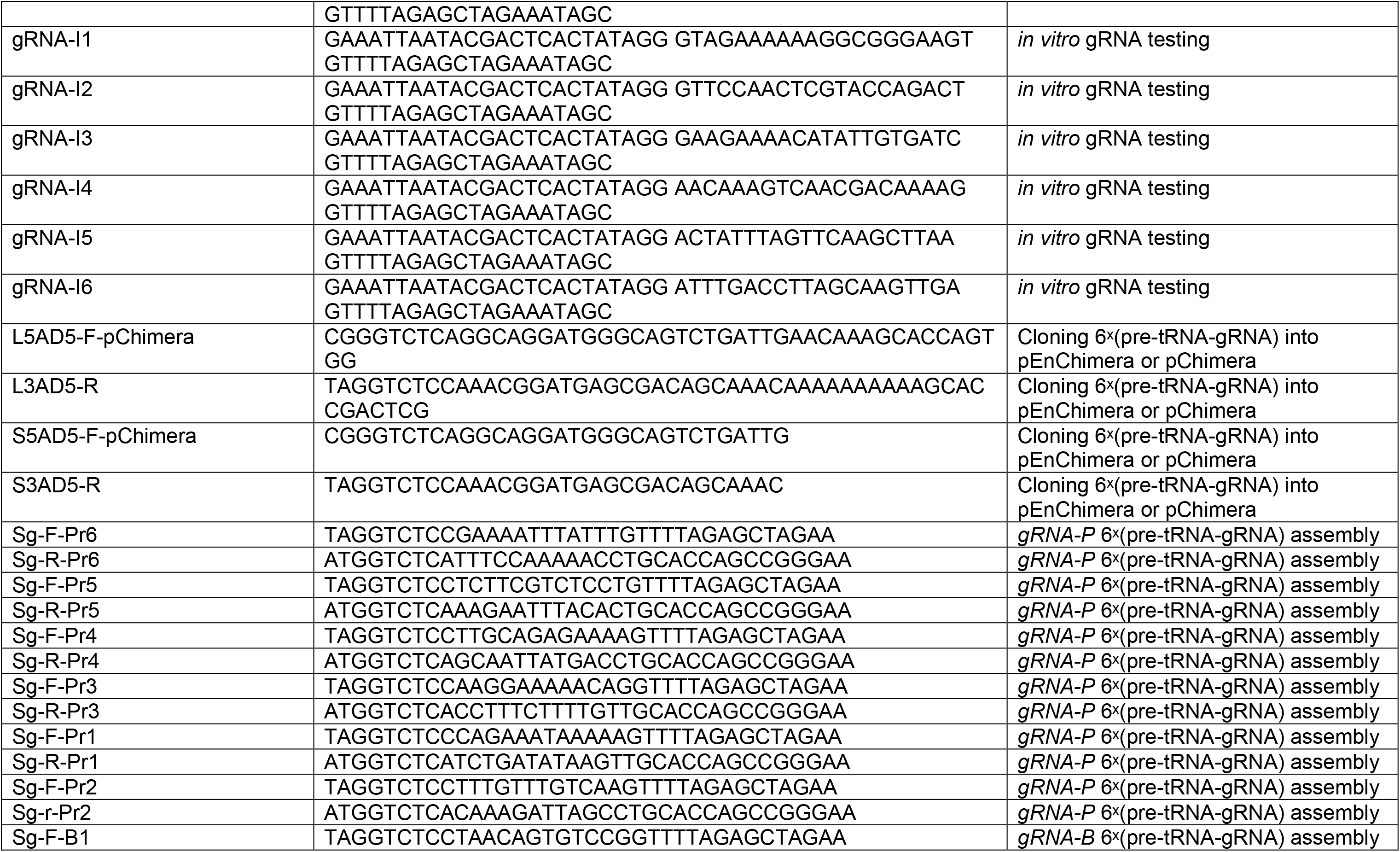
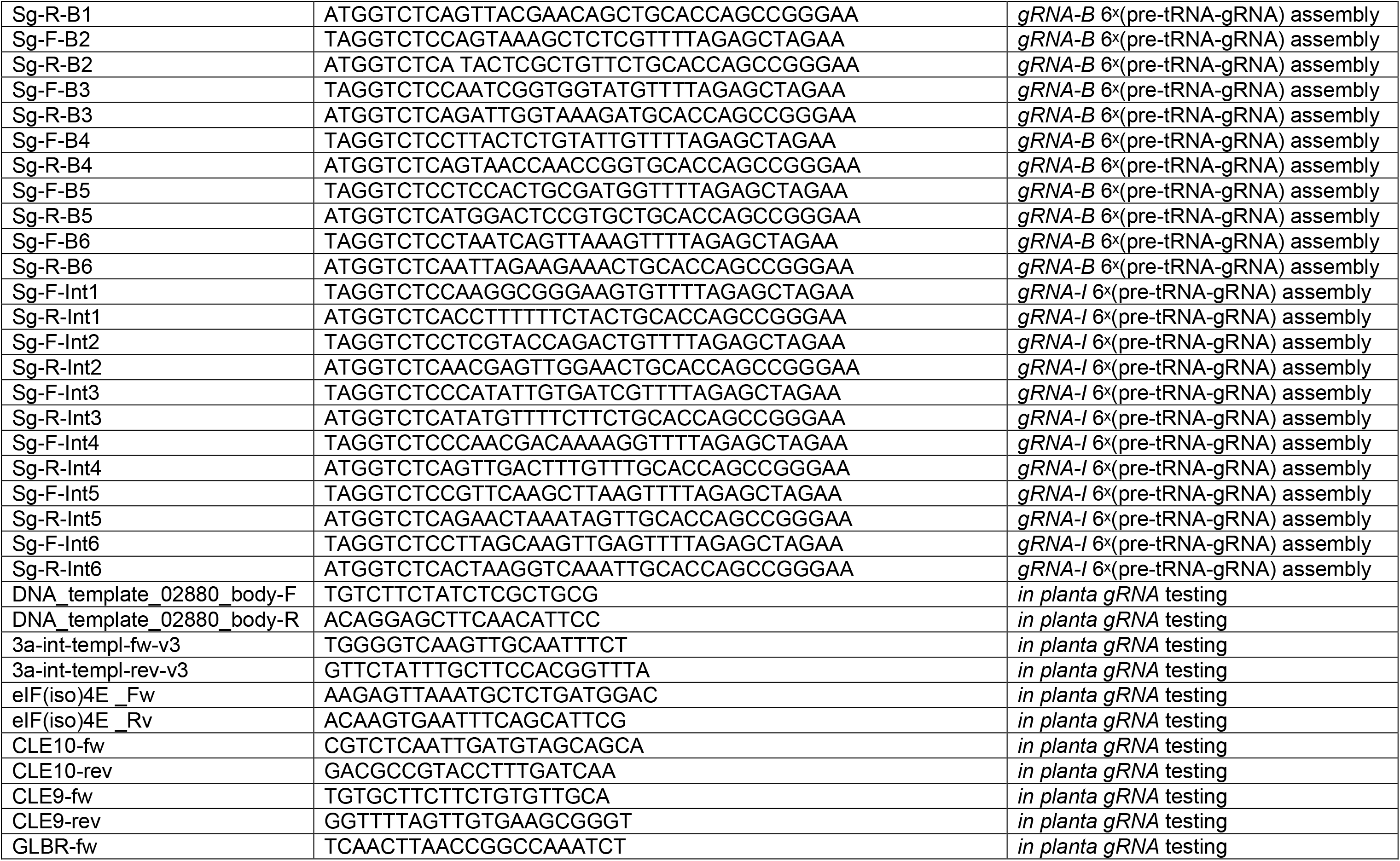
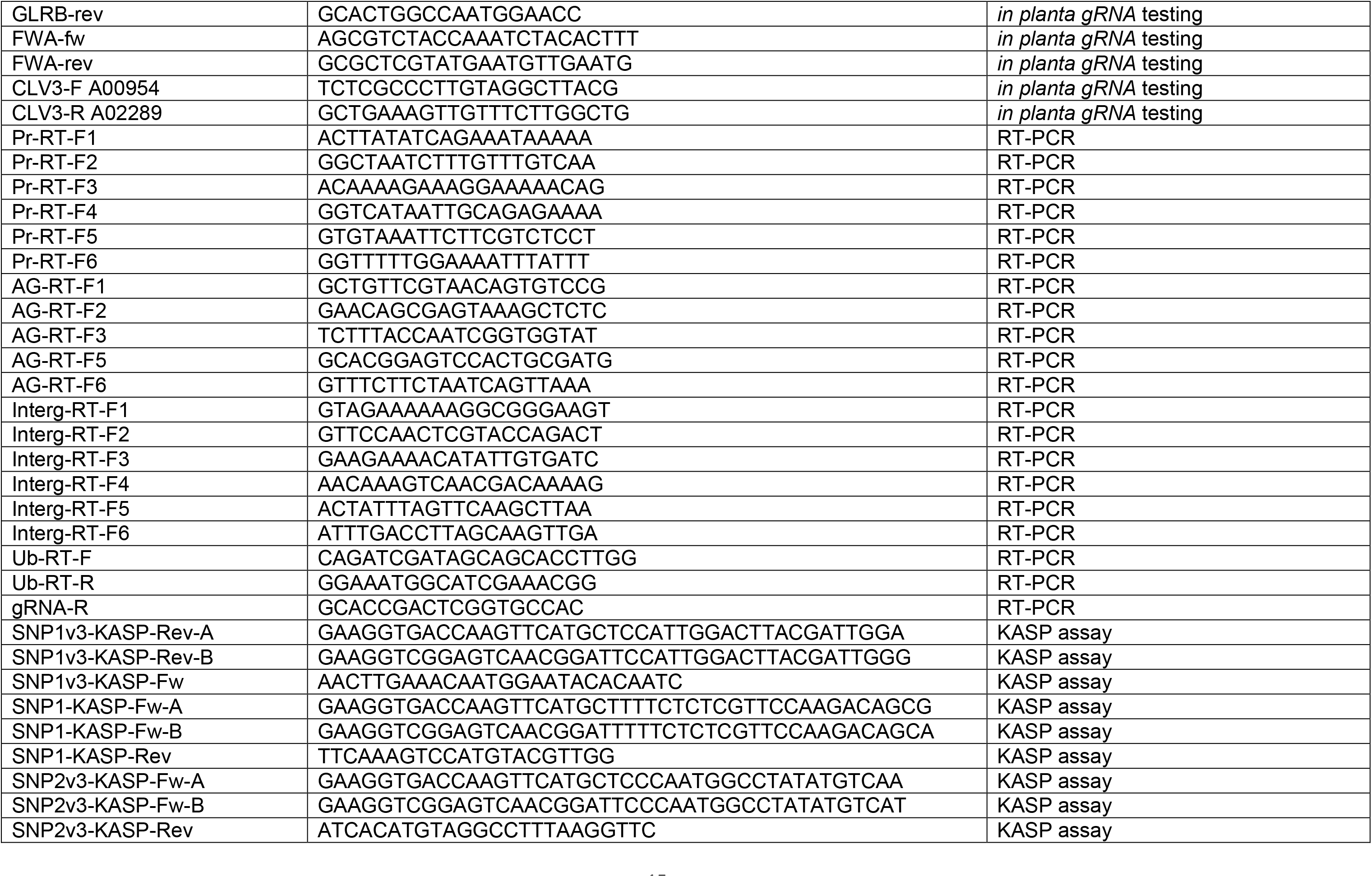
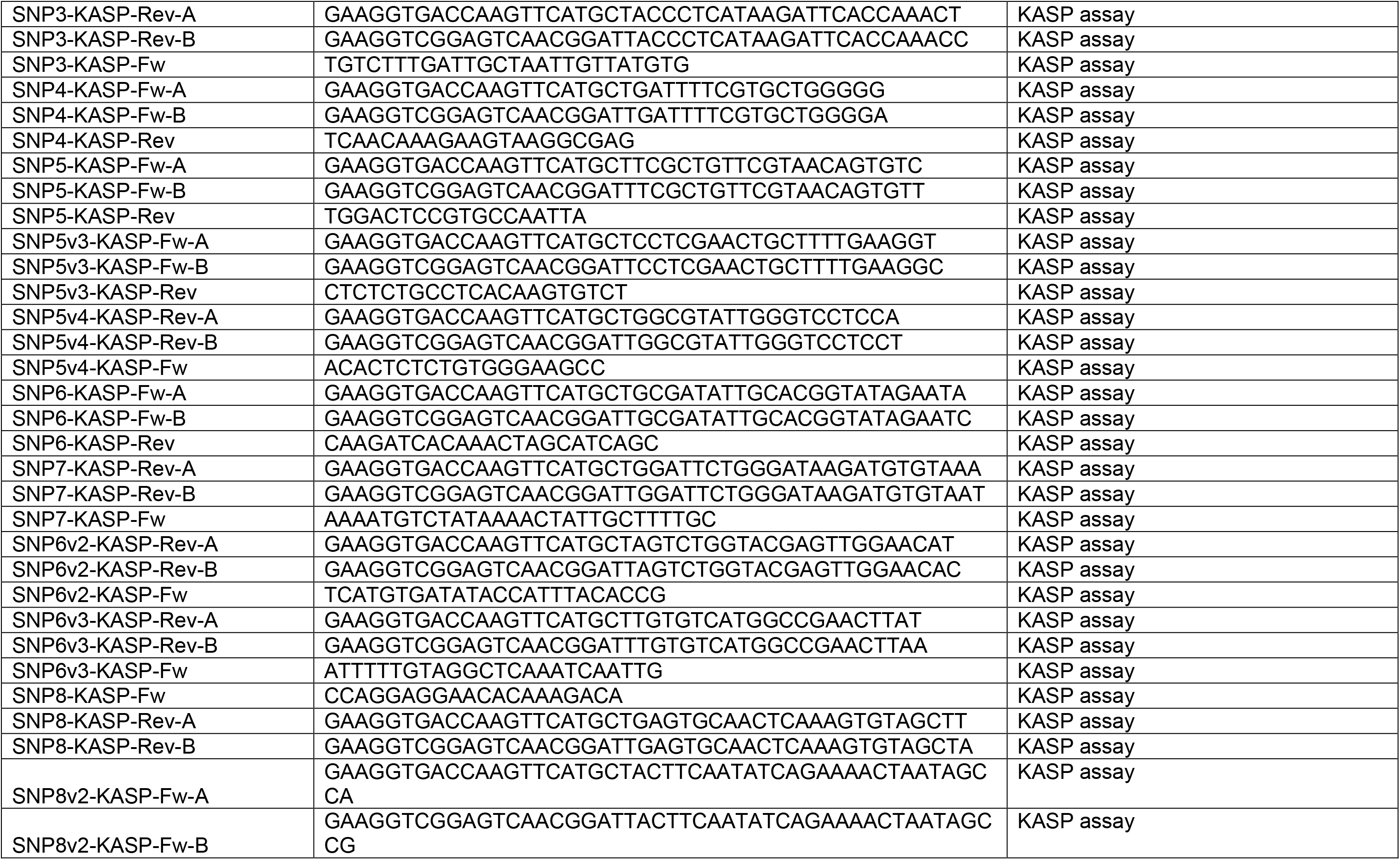
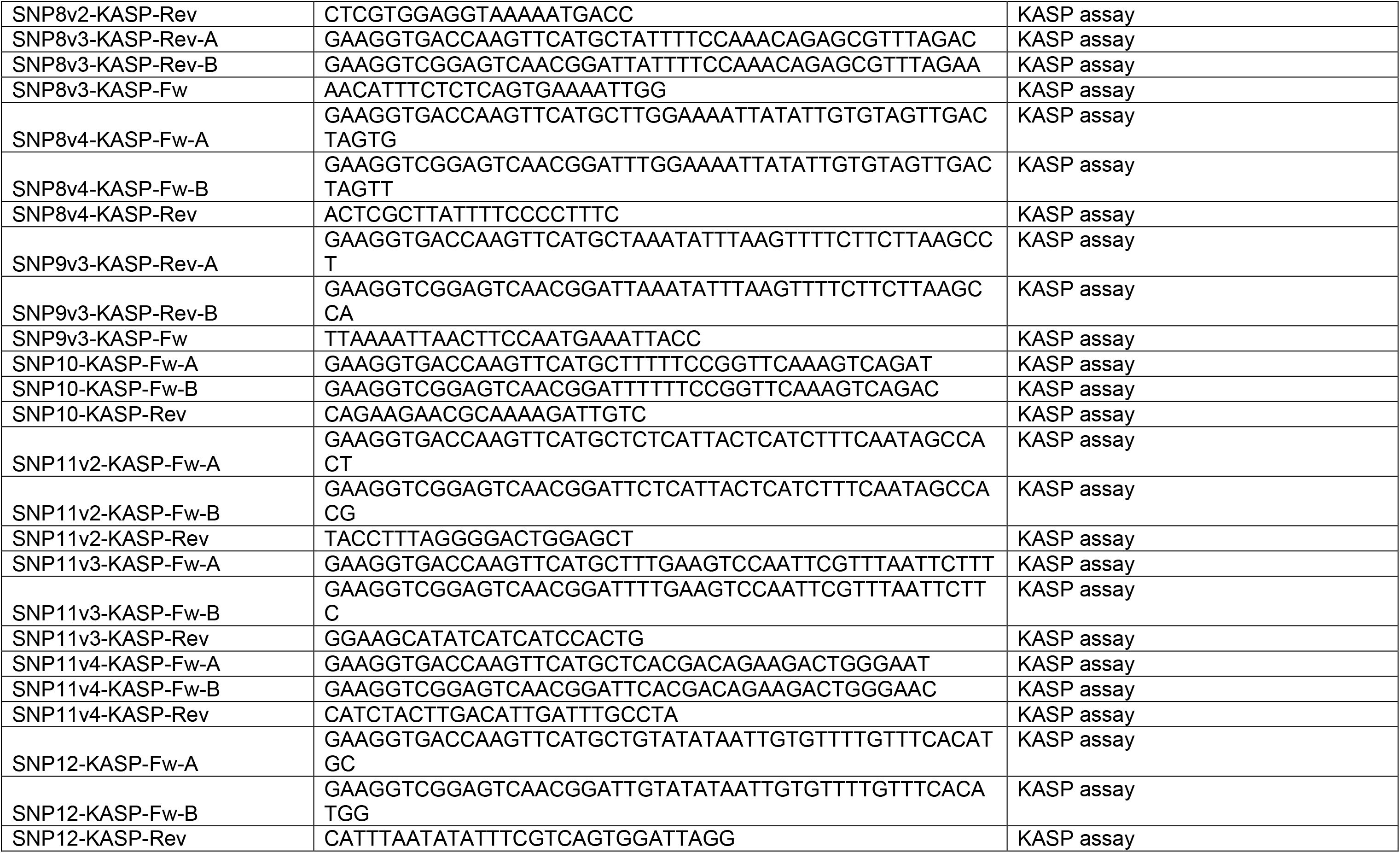
Oligonucleotides used in this study.

